# Multi-omics delineate growth factor network underlying exercise effects in an Alzheimer’s mouse model

**DOI:** 10.1101/2024.05.02.592289

**Authors:** Xin Li, Chaozhong Liu, Wenbo Li, Yanwan Dai, Chaohao Gu, Wenjun Zhou, Veronica C. Ciliberto, Jing Liang, Udhaya Kumar. S, Dongyin Guan, Zhaoyong Hu, Hui Zheng, Hu Chen, Zhandong Liu, Ying-Wooi Wan, Zheng Sun

## Abstract

Physical exercise represents a primary defense against age-related cognitive decline and neurodegenerative disorders like Alzheimer’s disease (AD). To impartially investigate the underlying mechanisms, we conducted single-nucleus transcriptomic and chromatin accessibility analyses (snRNA-seq and ATAC-seq) on the hippocampus of mice carrying AD-linked NL-G-F mutations in the amyloid precursor protein gene (APP^NL-G-F^) following prolonged voluntary wheel-running exercise. Our study reveals that exercise mitigates amyloid-induced changes in both transcriptomic expression and chromatin accessibility through cell type-specific transcriptional regulatory networks. These networks converge on the activation of growth factor signaling pathways, particularly the epidermal growth factor receptor (EGFR) and insulin signaling, correlating with an increased proportion of immature dentate granule cells and oligodendrocytes. Notably, the beneficial effects of exercise on neurocognitive functions can be blocked by pharmacological inhibition of EGFR and the downstream phosphoinositide 3-kinases (PI3K). Furthermore, exercise leads to elevated levels of heparin-binding EGF (HB-EGF) in the blood, and intranasal administration of HB-EGF enhances memory function in sedentary APP^NL-G-F^ mice. These findings offer a panoramic delineation of cell type-specific hippocampal transcriptional networks activated by exercise and suggest EGF-related growth factor signaling as a druggable contributor to exercise-induced memory enhancement, thereby suggesting therapeutic avenues for combatting AD-related cognitive decline.

## INTRODUCTION

The beneficial effects of physical exercise on neurocognition are widely observed in human patients with Alzheimer’s disease (AD) and animal models^1^. The underlying mechanisms are not completely understood and likely multifaceted through a combination of metabolic, endocrine, immunological, and neuronal changes^2,3^. Transcriptional regulation is particularly interesting because it is a general component of signaling pathways elicited by metabolic, hormonal, or neuronal cues, which is implicated in neurodegenerative diseases such as AD^4^. In addition, transcriptional regulation has a relatively long-lasting effect compared to allosteric regulation or posttranslational modifications of proteins, which is temporally in keeping with the long-lasting effects of physical exercise on neurocognition^5^. Considering that the hippocampus controls memory and that memory dysfunction is a hallmark feature of cognitive decline in AD, we performed single-nucleus multi-omics analyses of the hippocampus to provide a panoramic view of the transcriptomic and chromatin accessibility responses to long-term physical exercise.

## METHODS

### Mice

C57BL/6J wild-type (WT) and APP^NL-G-F^ mice were housed in standard 12 h light/ 12 h dark conditions. For wheel-running, mice were put into cages with a running wheel. The sedentary control group was put into the same type of cages but with a locked wheel. Two mice were housed per cage to avoid social isolation except when mice must be separated and singly housed occasionally due to bullying or fighting. The wheel-running activity was monitored in real time by the Actimetrics ClockLab data collection system. Gefitinib (AdooQ Bioscience A10422) and Wortmannin (AdooQ Bioscience A11161) were dissolved in DMSO and diluted in saline, followed by oral gavage at 50 mg/kg and 0.5 mg/kg, respectively, once every other day. Recombinant mouse HB-EGF protein (Novus NBP2-35069) was dissolved in saline and administered intranasally in awake mice at 3 ug/mouse (around 100 ug/kg) with pipettor at about 3 ul per nostril with alternating rest periods and a total administration volume of 10 ul per day^6^. The sex, age, and duration of the running were indicated in figure legends for each experiment. Animal protocols were approved by the Institutional Animal Care & Use Committee (IACUC) at Baylor College of Medicine.

### Behavioral tests

All the behavior tests were carried out between 12 PM and 7 PM in a dim light environment (300 lumens) except the light-dark test. For the object-in-place test, the task comprised an acquisition and a test phase separated by a 24 h delay. Each mouse was habituated in an arena (40□cm□×□40□cm□×□30□cm) without objects for 5 min to minimize confounding anxiety and novelty factors. In the acquisition phase, each subject was placed in the center of the arena, which contained 4 different objects in the corners 10 cm away from the walls. Mice were allowed to investigate the objects for 5 min and returned to the home cage. All objects were cleaned with alcohol between each mouse. In the test phase, each subject was replaced in the arena with swapped positions for 2 objects, and the subjects were allowed to investigate the objects for 5 minutes. Swapped objects were randomly chosen for each mouse. The discrimination index was calculated as (total exploration time on the novel objects – total exploration time on the familiar objects) / (total exploration time on the novel objects□+□total exploration time on the familiar objects). Considering that normal aging is probably associated with only a mild cognitive decline, we used a modified object in-place test to increase the difficulty of the test and, therefore, to expose the cognitive difference between the exercise group and the control group in the wild-type mice^7^. Briefly, the mouse was allowed to freely explore for 10 min in an arena with 5 Lego-built objects of different shapes and colors (sample phase). After 24 h, two of the objects were relocated. The mouse was re-introduced to the arena and was allowed to explore for 5 min (choice phase). Mouse behaviors were videotaped, and the discrimination index in the choice phase was calculated as the ratio of the time spent exploring the objects with the new location versus the total time spent exploring any object.

Novel object recognition (NOR) test was performed as previously described ^8^. Briefly, mice were habituated in an arena (22 cm x 44 cm) for 5 min. During the training session, two identical objects built from Lego were placed on the right and left sides of the arena. During the first day, mice were placed in the center of the arena and allowed to explore freely for 5□min. Mice were returned to their home cages. After 24 h, on the test day, one of the objects was replaced by a novel object (built from Lego) with a different color and shape, and mice were allowed to explore the arena for 5□min. Animal behavior during the training and test session was tracked by a top camera and analyzed by ANY-maze software (Stoelting). The discrimination index was calculated as (exploration time on the novel object - exploration time on the familiar object) / (exploration time on the novel object□+□exploration time on the familiar object). Exploration behaviors were defined as sniffing or touching (>1□s) the objects while looking at the objects.

The Y-Maze short-term spatial memory was measured according to the established procedure ^9^. The Y-maze was made of white opaque acrylic and had three arms (height: 20 cm; length: 30 cm; width: 8 cm) 120 degrees apart. Spatial cues were placed on the walls in line with the axes of the familiar and novel arms. The paradigm consisted of a 5-minute encoding trial during which one arm was blocked off, followed by a 1-h intertrial interval, then a 5-minute retrieval session. The start arm remained the same in both the encoding and retrieval trials, while the exposed arm during the encoding trial was considered the familiar arm, and the blocked arm was considered the novel arm. The apparatus was wiped with 70% ethanol and dried between each mouse to remove odor cues. The discrimination index was calculated as the comparison between the times spent in the novel and familiar arms during the retrieval paradigm.

The social interaction and social memory test was performed in a three-chamber apparatus (40.5cm x 60 cm x 22 cm) that had three chambers (left, center, and right) of equal size with 10□×□5□cm openings between the chambers. Mice were given 5 min habituation in the chamber and two consecutive 10□min tests: the first test measured sociability by subjecting the mouse to an intruder under one mesh pencil cup and an empty pencil cup, and the second test measured social novelty by subjecting mice to a novel intruder under the empty pencil cup. A camera and the ANY-maze software program were used to track the mouse in the three-chambered box while the experimenter scored the approaches to the object or partner mouse using a wireless keyboard. Intruders (sex-, age- and weight-matched) habituated to the mesh pencil cups in the apparatus for 1□h per day for 2 days before testing. Intruder mice were used up to three times, with one test per day.

The Morris water maze (MWM) test was performed as described previously ^8^. The MWM was virtually divided into four quadrants. During the training session, a transparent rescue platform was submerged under the painted water (0.5 cm –1 cm) and placed in a fixed position between the south and east quadrants of the pool. On the first day of training, mice were first allowed to stand on the platform for 10□s. After that, mice were gently placed into the water facing the wall and allowed to explore for 1□min. Mice were then guided to the rescue platform if they did not find it. Mice were allowed to take a rest on the platform for 10□s, and then re-trained from a different start position with the same procedure. After four training trials, they were dried using a paper towel and returned to home cages. Twenty-four hours later, mice were trained again following the same procedure without the initial habituation session. Mice were trained for five consecutive days. At the end of the fourth trial on day 5, mice were returned to home cages for a rest. One hour later, mice were put into the water maze from the west quadrant and let the mice explore the water maze for 1□min, where the platform had been removed. Mouse behaviors were videotaped and analyzed by the Noldus EthoVision XT. The rescue platform was located in the target quadrant. Mouse memory is evaluated by the escape latency and percentage of time spent in the target quadrant. Escape latency was defined as the time spent before finding the platform. Escape latency during the 5-day training sessions served as an independent measurement of spatial learning and memory.

The open-field arena (OPA) test was performed using the Versamax animal activity monitor equipped with infrared photo beams as horizontal *X*-*Y* sensors and/or *Z* sensors. Mice were placed in the center of the open-field arena (40□cm□×□40□cm□×□30□cm) and allowed to explore for 40□min. The locomotor activity and location of the mice were scored automatically by VersaMax software. The percentage of time spent in the center area measures anxiety levels. For the elevated plus maze (EPM) test, we used a plus-shaped platform that was elevated to 40□cm above the floor. Two opposite arms of the maze were walled (15□cm high), whereas the other two arms were open with a 5□mm high ridge to prevent falling. Each arm was 8□cm wide and 25□cm long. The test lasted for 10□min and was started by placing a mouse in the center part of the maze, facing one of the two open arms under a dim environment (300 lumens). An overhead camera and the ANY-maze software program were used to track the mouse. The time spent in the open arms was used as a measure for anxiety. The light-dark (LD) test was performed in a box developed from the open field chamber by placing a dark chamber occupying one-third of the open field box. The light area was connected by a small opening to allow mice to move from one area to the other. The test lasted for 10□min in a bright environment and started by placing a mouse in the bright area. The activity and location of the mouse were scored automatically by VersaMax software. The number of transitions between dark and light zones and the time spent in the light and dark areas were the index for anxiety.

### Histology, RNAscope, and ELISA

Mice were anesthetized with isoflurane (3-4% for induction, 1.5-2.5% for maintenance) for transcardiac perfusion with cold PBS and 4% paraformaldehyde. Overnight-fixed brains were immersed in 30% sucrose, embedded in the optimal cutting temperature (OCT) compound, and frozen in isopentane in dry ice. Coronal brain sections (30□µm) were prepared on the Leica CM1850 cryostat slicer. The coronal sections were collected in cryoprotectant solution (25% glycerol, 25% ethylene glycol, 50% PBS pH 7.4). Anti-β-Amyloid (6E10) antibodies (Biolegend, 803001;1:500), Fluor 488 goat anti-mouse IgG(H+L) (Life Technologies, A11029; 1:1,000) were diluted in TBS blocking buffer separately before use. Antigen retrieval was performed for 6E10 antibody by formic acid treatment (90%formic acid for 5 min for 5 min at room temperature (25 °C)). Brain sections were incubated with the primary antibodies at 4 °C overnight. Sections were then washed three times in TBS at room temperature and incubated further with fluorescence-conjugated secondary antibodies for 1□h at room temperature. These sections were then washed three times in TBS at room temperature, mounted with DAPI Fluoromount-G (SouthernBiotech, 0100-20), and sealed with the coverslip. Immunofluorescence of brain sections was viewed and captured with the Zeiss Axio imager.M2m microscope (Axiovision 4.8) and processed by ImageJ software (v 1.53e). The immunoreactive areas were quantified using ImageJ as previously described^10^. The average data of at least three sections per mouse was used to reduce the variance among tissue sections. A two-way ANOVA followed by a post-hoc Fisher’s LSD test was used to analyze the quantification data.

For RNAscope analysis, mice were anesthetized for transcardiac perfusion with cold PBS and 4% paraformaldehyde. Overnight post-fixed brains were immersed in 30% sucrose, embedded in OCT, and frozen in precooled isopentane. Coronal brain sections (around 12 μm) were prepared on the Leica CM1850 cryostat slicer. The coronal sections were collected. Stxbp1 was assayed using RNAscope following the standard protocol from ACD with minor modifications. In brief, brain sections were rinsed with PBS to remove OCT. The brain sections were incubated at 60□°C for 30 min. Then, the brain sections were post-fixed in 4% PFA at 4□°C for 15 min. After the post-fixation, the brain sections were dried in ethanol. The brain sections were then incubated with hydrogen peroxide at room temperature for 10 min. The sections were rinsed for 2 min three times in distilled water, and then the brain sections were retrieved in RNAscope 1× target retrieval reagent at 100□°C for 5 min. The slides were then rinsed in distilled water for 2 min three times and re-dried in 100% alcohol for storage. The pretreated brain sections were incubated with protease III for 30 min at 40□°C. The protease III was removed, and the brain sections were rinsed in distilled water for 2 min three times. The brain sections were hybridized with the probe of Stxbp1 (ACD) for 2 h at 40□°C. After that, the brain sections were rinsed for 2 min three times in the wash buffer to remove the excessive probes. The RNAscope Multiplex FL v2 Amp1 was added to the brain sections and incubated at 40□°C to amplify the signal for one probe. The brain sections were rinsed with wash buffer after 30 min. The probe signals were detected using the RNAscope Multiplex Fluorescent Detection Reagents V2 (ACD 323110). Brain sections were treated by the Multiplex FLV2 HRP blocker and washed in PBS for 2 times. Brain sections are viewed and captured with the Zeiss Axio imager M2m microscope and processed by ImageJ software. To quantify the expression of Stxbp1, we measured the counts of the signal dots in each cell within the specific regions using ImageJ software. At least 10 neurons were counted for each mouse, and the averaged number from those cells represents the expression intensity for each mouse. The results were calculated as counts per cell. A student’s two-tailed t-test was used to analyze the quantification data.

Mouse serum HB-EGF levels were determined by the Mouse HB-EGF ELISA kit (ABclonal, RK02882). Blood was collected through the tail vein using a microvette capillary blood collection tube. Samples were incubated at room temperature for 40 min, and centrifuged at 3,000 g for 15 min to collect serum. ELISA is performed according to the manufacturer’s instructions.

### snRNA-seq and snATAC-seq

Mice were euthanized with CO_2_ after wheel-running for 3 months at the age of 6 months old. The hippocampus was isolated immediately, washed with cold D-PBS, and snap-frozen with liquid nitrogen. Hippocampi from 3 mice were pooled together before nuclei isolation and capture for each of the analyses (snRNA-seq and snATAC-seq). The nuclei isolation for snRNA-seq and snATAC-seq were performed in different batches from different mouse cohorts. The Frankenstein protocol was adopted to isolate nuclei from frozen tissues (dx.doi.org/10.17504/protocols.io.3fkgjkw). Briefly, the hippocampus was sliced into 2-5 mm^3^ pieces on ice, and the samples were transferred to a 1 ml Dounce homogenizer with prechilled nuclei isolation buffer (10 mM Tris-HCl, 10 mM NaCl, 3 mM MgCl_2_, and 0.01% NP40). The tissue was homogenized with five strokes of the loose pestle and 10 strokes of the tight pestle. The homogenate was filtered through a BD Falcon 40 µm cell strainer and centrifuged at 300g for 5 minutes at 4 °C. The supernatant was discarded. The nuclei pellet was resuspended with D-PBS with 20% percoll (GE) and centrifuged at 500 g for 15 minutes at 4 °C. The supernatant was discarded, and the nuclei pellet was resuspended D-PBS with 1% BSA for snRNA-seq and with the Nuclei Buffer for snATAC-seq. Nuclei concentration was measured with a hemocytometer, and the samples were further processed using the NovaSeq system at the Single Cell Genomics Core at Baylor College of Medicine for 10x single cell 3’ v3 RNAseq and 10x Single Cell ATAC sequencing. An average of 10k cells were targeted for capture per sample per assay, with over 50,000 reads per cell. snRNA-seq and snATAC-seq were performed separately on different cohorts of mice, generating 246,765,198 reads from 34,950 nuclei for snRNA-seq and 211,976,010 reads from 31,793 nuclei for snATAC-seq after quality control.

### snRNA-seq data initial processing

Sequenced reads were first processed and quality controlled using the Cell Ranger Single Cell Software Suite provided by 10x Genomics (version 3.1.0). Reads were aligned to the mouse genome (mm10), and read counts per gene for every cell in a sample were obtained. We used CellBender (version 0.1.0, https://github.com/broadinstitute/CellBender.git) to identify nuclei with ambient RNA further using the remove-background function with the ‘total-droplets-included’ set at 1,000,000. Nuclei kept by both Cell Ranger and CellBender were retained for subsequent analyses. A total of 37303 nuclei were included (8804 APP_EX, 10411 APP_RT, 6891 WT_EX, 11197 WT_RT). Seurat pipeline was used to analyze the kept nuclei by first removing cells with more than 5% of reads mapped to mitochondrial genes and those with less than 500 genes or more than 5500 genes. On the remaining 32285 cells, the top 2000 variable genes were identified and used to perform dimension reduction using PCA with a maximum of 200 PCs. The first 50 PCs were then used for further clustering and UMAP. To decide the cell populations, we used clustering with a resolution of 0.5. The top marker genes were identified empirically using the FindMarkers function. To identify differentially expressed genes (DEG) between the genotypes in each cell population, we first subset the cells of the particular cell population and then used the FindMarkers function from Seurat.

Gene Set Enrichment Analysis (GSEA) v4.2.2 Mac App was used. Two libraries specific to mouse, mouse pathway (893 gene sets) and GO libraries (12526 gene sets) were obtained from http://ge-lab.org/gskb/; along with C2 (6290 gene sets) and C5 (14998 gene sets) from MSigDB (https://www.gsea-msigdb.org/gsea/msigdb/index.jsp) were used in the analysis. GSEPreparked analysis was adopted with the ranked files generated from logFC of all genes except for the ones with no expression in each cell type obtained from the FindMarkers function from the Seurat package. The logFC was calculated with SCTransformed data in the Seurat object. GSEA was run for each cell type in three contrasts: APP_EX vs. APP_RT, APP_RT vs. WT_RT, and WT_EX vs. WT_RT. Parameters used in the GSEA were: max size = 5,000; min size = 5; collapsing mode for probe sets => 1 gene = max_probe; normalization mode = meandiv. The GSEA analysis results were examined at different thresholds: 1. Soft threshold with p < 0.001; 2. Median threshold with FDR < 0.25 and p < 0.001; 3. Harsh threshold with FDR < 0.1 and p < 0.001; 4. Extreme threshold top 10 in normalized enrichment score (NES) (positive/negative) and FDR < 0.25 and p < 0.001. Three gene sets were shortlisted for further analysis. They were picked out of all gene sets that were significantly over-represented at the median threshold in more than 4 cell types and were reversed in NES in APP_EX vs. APP_RT and APP_RT vs. WT_RT contrasts.

Sub-clustering analysis was done in excitatory neuron (EN) nuclei using Seurat version 3.2.2. No further filters were applied for nuclei in the sub-clustering analysis. The SCT assay matrix of all excitatory neuron nuclei was used. Variable features were identified within the chosen cell type’s nuclei with the function FindVariableFeatures with the vst method, and the top 2,000 features were used in the analysis. Then, dimensionality reduction was run using the RunPCA function. Data was visualized by UMAP with dim 1:30. Then EN nuclei were clustered with the function FindNeighbors with dims 1:30 and FindClusters at resolution 0.2. Sub-cluster markers curated from literature^11^.

### snATAC-seq data initial processing

The data from sorted bam files created by CellRanger (version 3.1.0) was processed with package snaptools to generate the bin-by-nuclei matrix for ADEX, ADRT, WTEX, and WTRT samples separately, following the online tutorial: https://github.com/r3fang/SnapTools. Bam files were processed by the snap-pre function with parameters settings as min-mapq = 30, min-flen = 50, max-flen = 1000, keep-chrm = TRUE, keep-single = FALSE, keep-secondary = False, overwrite = True, num = 20000, min-cov = 500. Then snap-add-bmat function was used to call the bin matrix with bin-size-list set as 5,000. All samples were then combined for analysis with the package snapATAC following the pipeline tutorial https://github.com/r3fang/SnapATAC/blob/master/examples/10X_brain_5k/README.md. Original filters of nuclei were set as UMI >= 3 & <= 5; promoter_ratio >= 0.15 & <= 0.6; peak_region_fragments > 1,000 & < 20,000; frag_in_peak_ratio > 0.15. Bins in the blacklist or mitochondrial chromatin or over top 95% coverage among all bins were filtered out. Dimensionality reduction was done with the function runDiffusionMaps with num.eig set as 50. Clustering was done with the function runKNN (eigs.dims set as 1:30 and k as 15) and the function runCluster (leiden version 0.3.5 and resolution as 1.5). Bins were binarized for further analysis. Peaks were then called on bins in each cluster with buffer.size set as 500. Peaks were then merged by the function reduce. Lastly, peaks were scaled by the function scaleCountMatrix with the RPM method. Snaptools and SnapATAC are installed from GitHub: https://github.com/r3fang/SnapATAC (Snaptools version v1.2.3 and SnapATAC version 1.0.0).

The peak-by-nuclei matrix was then imported to the Seurat pipeline for further analysis. Peaks from the blacklist, unwanted chromosomes, and small clusters (Cluster 18, 19, and 20) were filtered out. Furthermore, nuclei with less than 1,000 peaks were removed. UMAP was run again in Seurat to generate the embedding for visualization. ACTIVITY matrix for gene-by-nuclei matrix was generated by summing up peaks within 2kb upstream of the TSS (transcription starting site). Based on the ACTIVITY matrix, marker plots and enrichment analysis were carried out to determine the final cell type annotations. Sub-clustering analysis was done in excitatory neuron nuclei using Seurat version 3.2.2. No further filters were applied for nuclei in the sub-clustering analysis. The ACTIVITY matrix of all excitatory neuron nuclei was used. Variable features were identified within the chosen cell type’s nuclei with the function FindVariableFeatures with the vst method, and the top 30,000 features were used in the analysis. Then, dimensionality reduction was run using the RunLSI function. Data was visualized by UMAP with dim 1:30. Then EN nuclei were clustered with the function FindNeighbors with dims 1:30 and FindClusters at resolution 0.2. Sub-cluster markers^11^.

### Integrated multi-omics data processing

The snRNA-seq and snATAC-seq data were integrated following the Seurat integration pipeline. The assay used for snRNA-seq was RNA, and for snATAC-seq, it was ACTIVITY. Anchors were found with the function FindTransferAnchors with variable genes from RNA data as features, reference.assay as “RNA”, query.assay as “ACTIVITY”, reduction method as cca, and dims with 1:40. ATAC data was then imputated with the function TransferData using anchors found, weight reduction with LSI method, and variable genes as variable features from RNA object. Then snATAC and snRNA data were merged with the function merge. RunPCA was performed on integrated data with all features in the combined data and RunUMAP with dim 1:30.

Trajectory analysis was done in oligodendrocytes following the monocle3 trajectory analysis pipeline (monocle3 version 1.0.0). Oligodendrocyte nuclei were sub-clustered into two sub-clusters with FindNeighbors (dims 1:30) and FindClusters (resolution 0.05). The bigger sub-cluster with 10,480 nuclei out of all 11,252 nuclei was used in trajectory analysis. The learn_graph and order_cells were applied with default monocle3 settings. Calculated pseudotime was then binned in equal 50 frames for analysis. In plotting, each bin on the x-axis was represented by the middle value of each binned frame. Proportions for nuclei within each bin were calculated by the formula shown below:

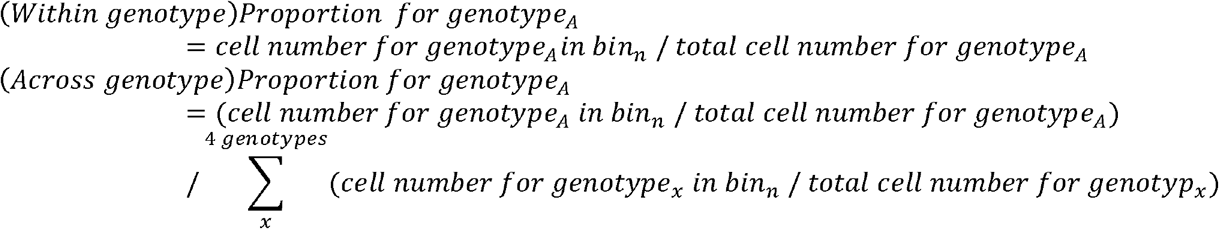

Module scores were calculated by the Seurat function AddModuleScore for nuclei in each of the 50 binned pseudotimes. Oligodendrocyte marker lists are from literature ^12^. Newly formed oligodendrocyte markers: Tcf7l2, Casr, Cemip2, Itpr2; Myelin forming oligodendrocyte markers: Mal, Mog, Plp1, Opalin, Serinc5, Ctps1; Mature oligodendrocyte: Klk6, Apod, Slc5a11, Pde1a.

### Differential Accessible Regions detection by a Siamese neural network model

Due to the lack of power in using statistical tests to discover DARs, we instead trained a simple neural network to distinguish different conditions, followed by interpreting the trained model to extract the differentially accessible regions. We detected DARs for each cell type separately. Preprocessed and cell-type annotated snRNA-seq and snATAC-seq were loaded as Seurat v3 objects. Then, cell-type-specific snRNA-seq and snATAC-seq Seurat objects were created to perform omics data integration by Seurat v3. After integration, anchor cell pairs inferred by Seurat were saved, together with a highly-variable gene matrix (vst method, 5000 genes) and highly-accessible peaks matrix (detected in more than 5% of the cells). Gene matrix, peak matrix, anchor pairs, and genotype label were provided to train a customized Siamese neural network (explained in the next section). After training, the network was able to project all cells from different omics data into a common low-dimensional space, where snRNA-seq and snATAC-seq data are mixed, and different genotypes are separated from each other. With the goal of separating genotypes achieved, we believe the neural network has identified peaks that vary among genotypes. To extract these peaks from the black box model learned, we first applied the Activation Maximization algorithm (explained later) to construct 12 pseudo-cell snATAC-seq data which are identified by the model with a high possibility of belonging to each genotype. Then t-test was applied to find the peaks that are important exclusively in only one genotype. Directions of these peaks in APP_RT vs. WT_RT and APP_EX vs. APP_RT were determined by checking our real snATAC-seq data. Finally, peaks with log2 foldchange smaller than log2(1.1) or not falling in gene regions were removed. With all the above being done, we got a set of differential accessible regions among different genotypes in each cell type. In detecting DARs in this way, we believe it can bear the noises of single-cell data and utilize the information from snRNA-seq data. The regions found by this method were validated in the original data.

### Customized Siamese neural network to separate multi-omics data by genotypes

The goal of this neural network is to separate genotypes while mixing snRNA-seq and snATAC-seq data. After training, we will be able to find important peaks that can separate genotypes not only based on chromatin accessibility but also transcriptome. To do this, we created a model based on scDGN^13^ neural network framework, including several modules.

The encoder module is used to project datasets into a common lower dimensional space and contains two fully connected layers that produce the hidden features *x*′ = *f*_*e2*_ (*f*_*e*1*x*_(*x*; *θ*_ *e*1*x*_); *θ*_*e*2_ for snRNA-seq or *y*′ = *f*_*e2*_ (*f*_*e*1*y*_(*y*; *θ*_ *e*1*y*_); *θ*_*e*2_ for snATAC-seq, where *θ* represents the parameters in these layers. The label classifier, *f*_*lx*_ (*x*′; *θ*_*lx*_)and *f*_*lx*_ (*y*′; *θ*_*ly*_), ensures genotypes are separated in the common space. The goal of the domain discriminator *f*_*d*_ (*x*′; *θ*_*d*_) and *f*_*d*_ (*y*′; *θ*_*d*_) is to determine whether a pair of inputs ((xi,xj), (xi,yj), (yi,xj), (yi,yj)) are from the same domain or not. The overall objective function to be minimized is:

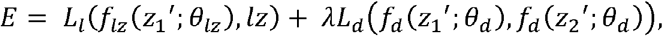

where *z*_1_,*z*_2_ ∈ {*x*,*y*}, *λ* can control the trade-off between the goals of domain invariance and higher classification accuracy. Inspired by Siamese networks, the domain loss adopts a contrastive loss for a pair ***z***_**1**_ and ***z***_**2**_, where ***z*** ∈ {*x*,*y*} :

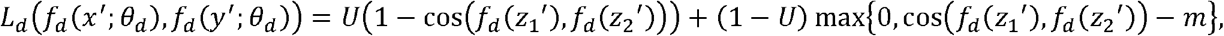

where U=0 indicates the two cells are from the same modality, but different genotypes, and U=1 indicates that they are identified as anchors by Seurat. *cos*(·) is the cosine embedding loss, and m is the margin that indicates the prediction boundary. Overall, the aim of the objective function is to minimize the label classification loss and the domain loss. In this way, genotypes are separated, and important peaks are learned by the model based on not only snATAC-seq but also snRNA-seq data.

### Activation Maximization to find important peaks

With genotypes separated, we used activation maximization to extract important peaks for each genotype. Given a particular genotype i and a trained neural network *f*, activation maximization looks for important input genes *x*_*m*_ and peaks *y*_*m*_ by solving the following optimization problem:

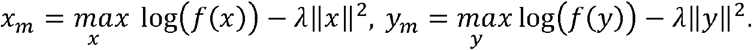

Twelve such pseudo-cells were constructed for each of the 3 genotypes, including WT_RT, APP_RT, and APP_EX. A t-test was performed to identify exclusively important peaks in genotypes, and their log-fold-change is determined by our snATAC-seq data.

### Homer motif analysis and regulatory network construction

With exercise reversed DARs of each cell type, we ran Homer^14^ *findMotifsGenome* function to perform known motif enrichment analysis. All parameters are set to default. The summary plot containing the p-value and peak ratio in each cell type, and reverse direction is drawn with ggplot2 in R. With discovered motifs in each cell type, we collected their downstream genes from TRASFAC database. Network per cell type was constructed, with edges indicating TF-downstream gene relationship. The networks were saved as .gml files and visualized in Cytoscape with color indicating log fold changes in the two comparisons.

### Integrated analysis with spatial transcriptomics data

The count matrix of spatial transcriptomics data was filtered by removing spots with tissue coverage of less than 30% in the HE images and then removing genes that were detected in less than 10 spots. The edgeR function “cpm” was used for the normalization of the filtered matrix. The output log cpm matrix was used for the following analysis. For assigning cells in the snRNA-seq data back to the locations in the spatial transcriptomics data, we suppose that the reference atlas has n positions with p genes, and the snRNA-seq data set has m cells with the same number of p genes. We aimed to assign the m cells into n positions using a linear regression model with L1 norm and generalized L2 norm via graph Laplacian. We created a random walk normalized graph Laplacian matrix based on the location information and anatomical structures from the ST data. If two spots belong to one anatomical structure and the distance between the spots is smaller than a specific threshold, then the spots will be connected in the Laplacian matrix. Our model uses a linear method to measure the differences in gene expression levels in assigning cells to locations. The optimal solution minimizes the differences between gene expression levels of individual cells and gene expression levels of locations. For each individual cell, we want to minimize the following objective function,

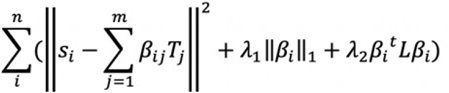

where *s ∈R* ^ *n × g*^ is the single-nucleus expression matrix, *T* ∈ *R* ^ *m × g*^ is marker genes expression matrix of reference atlas (spatial transcriptomics data), L is normalized graph laplacian computed based on the location distance matrix and anatomical structures. The L1 norm penalization encourages sparsity on the coefficients, which guarantees that one cell can only be assigned to a small number of locations. The generalized L2 norm encourages the smoothness of the coefficients, which guarantees that cells with similar gene expression levels are more likely to be assigned to closer locations. To map the snRNA-seq data back to the mouse hippocampus, the reference profile was first established by using the spatial transcriptomics expression atlas of 500 highly variable genes at the 6-month age retrieved from Navarro et al. Reconstructed spatial expression patterns were validated by well-known genetic markers, such as Prox1 and Ociad2. Reconstructed patterns of marker genes were consistent with the FISH images from Allen Brain Atlas.

To reconstruct spatial expression patterns, we used the following steps: (1) Read the gene expression matrix from snRNA-seq and expression matrix from reference atlas (spatial transcriptomics data). (2) Construct the Laplacian matrix based on the location information. (3) Use CVX to solve the convex function with L1 norm and generalized L2 norm. (4) Assign cells to target locations based on the distribution of marker genes in the objective function. (5) Reconstruct the spatial patterns based on the expression profiles in the snRNA-seq data and cell locations from the mapped results. Fill the expression profiles from the snRNA-seq data in the assigned locations.

To calculate cell proportion in different anatomical regions, we first performed glm-SMA algorithm to assign the cells from snRNA-seq data back to the locations from the spatial transcriptomics data. Then, we counted the cell number in different anatomical regions and calculated the cell proportion of each cell type in DG and CA regions. To confirm the result was not generated by chance, we randomly shuffled the genotypes in the APP_EX and APP_WT samples and repeated the shuffling 100 times. Then, we recalculated the cell proportions and did the student t-test. Granule cell proportion increased in the DG region from AD_EX samples with a p-value < 2.2e-16.

### Statistical analysis

Statistical analyses were performed using SPSS (V.21.0, IBM) unless described otherwise in the above sections. No statistical methods were used to pre-determine sample sizes. Instead, sample sizes were determined based on previous publications for the relevant assays. Normality was tested by the Shapiro-Wilk test (n < 10) or D’Agostino-Pearson omnibus test (n > 10). For non-normal data or data with nonequivalent variances, the comparisons between two or multiple groups were tested with the Mann-Whitney test or the Kruskal-Wallis test, respectively. All tests were two-sided. All measurements were taken from distinct biological samples (mice or human subjects). Most comparisons between the two groups were analyzed using a two-sided, unpaired t-test. Body weight with multiple time points or Morris water maze tests were analyzed with repeated-measures ANOVA with Tukey’s post hoc test. For statistical significance, a two-tailed unpaired t-test, or one-way repeated ANOVA with Fisher’s LSD test multicomparisons, was used for experiments with two groups. The behavior test experimenter was blinded to the exercise or pharmacological treatment conditions during the early stage of analysis, such as counting the time duration from video clips. The statistical analysis and data plotting was then done by experimentalists who knew both genotype and treatment information. Animals were excluded and euthanized before behavior tests if they showed distress, infection, bleeding, or anorexia. All data were expressed as mean ± SEM. All data were individually plotted (Prism 9, GraphPad). The exact numbers of animals are reported in the figure legends. P < 0.05 is set as significance.

### Sex as a biological variable

We used both male and female mice and clarified the sex in the figure legends and methods sections. Most experiments were done in male mice, while female mice were also used. We did not find sex differences.

## Data access

The snATAC-seq (GSE237884) and snRNA-seq (GSE237885) data will be available at NCBI’s Gene Expression Omnibus (GEO) under the GSE237925 SuperSeries after this manuscript is officially published.

## Data availability

Data is available upon request to the corresponding authors.

## Code availability

Code is available upon request to the corresponding authors.

## RESULTS

### Exercise improves memory functions and induces cell-type-specific transcriptomic changes

We used the C57BL/6J homozygous knock-in mice containing the Swedish (NL), Beyreuther/Iberian (F), and Arctic (G) mutations in the gene for amyloid precursor protein (APP^NL-G-F^)^15,16^ as a model for AD because it lacks the artificial hyperactivity phenotype from APP overexpression, which resembles human AD pathophysiology^17,18^. For chronic physical exercise, we used voluntary wheel-running to minimize stress on mice. Wild-type (WT) and APP^NL-G-F^ mice were put in cages with running wheels at 3 months old. After exercise for around 6 months exercise, the exercise groups (WT_EX and APP_EX) shared similar wheel-running exercise volumes and lost similar amounts of body weight compared to their respective rest controls (WT_RT and APP_RT) (**Suppl Fig S1a-b**). The chronic exercise improved memory functions in APP^NL-G-F^ mice but not in WT mice in the classical object-in-place test (**Fig 1a-c**) and Y-maze test (**Fig 1d-f**) at the age of 10 months. Total travel distances in these tests remained similar between WT and APP^NL-G-F^ mice (**Suppl Fig S1c-d**). Although exercise appeared to have no effects in WT mice in these standard tests, a modified object-in-place test with 5 objects^7^ demonstrated a clear memory-enhancing effect of exercise in WT mice (**Fig 1g-i**). In summary, chronic wheel-running exercise improves learning and memory in both WT and APP^NL-G-F^ mice, with a more robust effect on the APP^NL-G-F^ mice.

**Figure 1.**
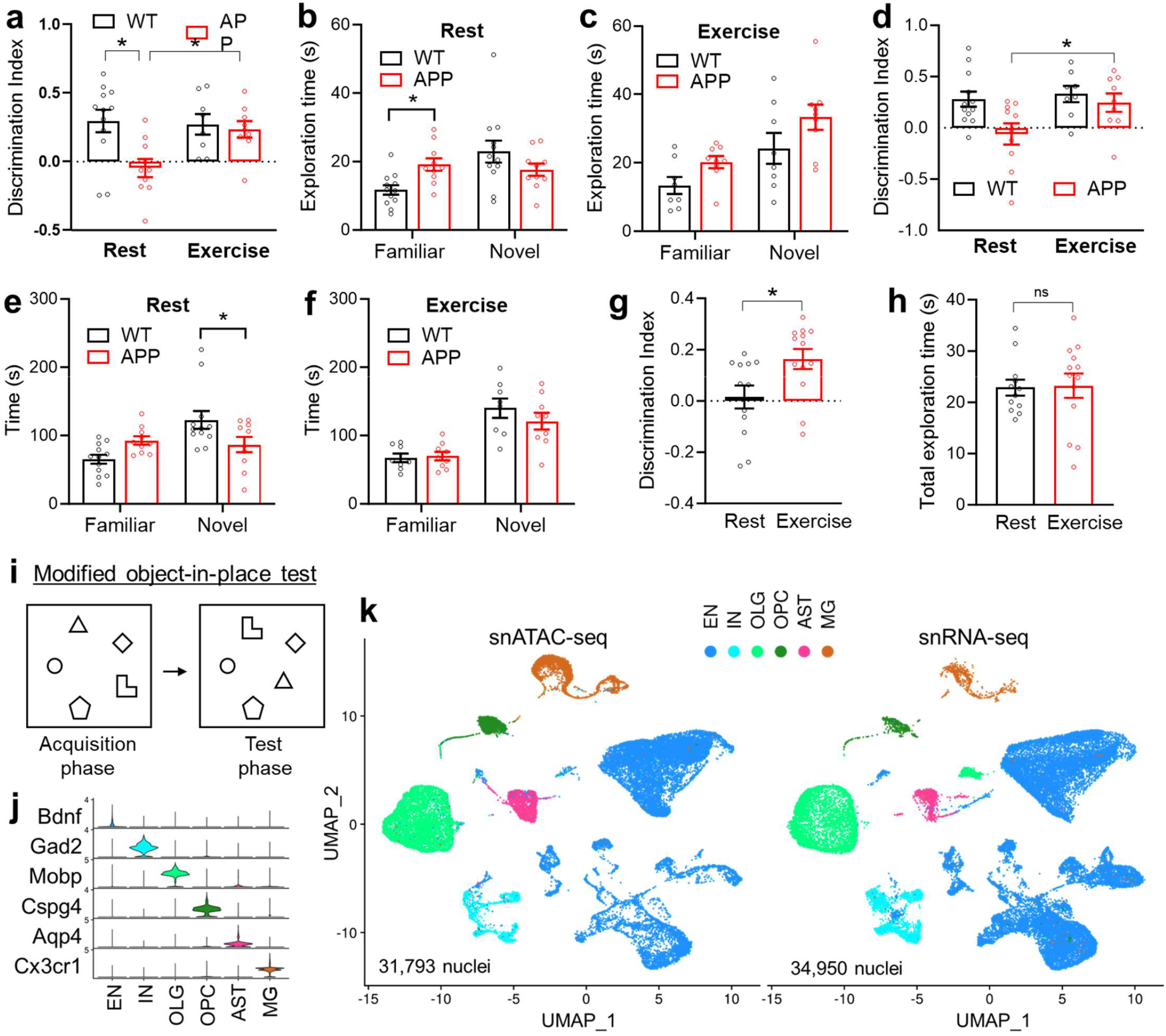
Exercise improves memory without altering the overall hippocampal cellular composition. Discrimination index and exploration time of the standard object-in-place memory test (**a-c**) and the Y-maze test (**d-f**) in male WT and APP^NL-G-F^ (APP) mice at 10 months old after 6 months wheel-running exercise (EX) or rest (RT) (n = 12 for WT_RT; 8 for WT_EX; 10 for APP_RT; and 9 for APP_EX). Asterisks indicate significant differences with the two-way ANOVA and Fisher’s LSD multiple comparisons test. (**g-i**) Discrimination index and exploration time of a modified 5-object in-place test in 4-month-old male WT mice after 2 months wheel-running (n = 12 for RT, 13 for EX). Bar graphs show the mean with S.E.M. Asterisks indicate significant differences by 2-sided t-test. (**j**) Violin plot of cell type-specific marker gene expression levels. (**k**) UMAP of major cell types of the hippocampus based on snRNA-seq and snATAC-seq datasets. Excitatory neurons (EN), inhibitory neurons (IN), microglia (MG), astrocytes (AST), oligodendrocytes (OLG), and oligodendrocyte progenitor cells (OPC). APP/Exercise (APP_EX), APP/Rest (APP_RT), WT/Exercise (WT_EX), WT/Rest (WT_RT).

We reason that the molecular changes in the brain would precede the behavioral changes and exercise for too long might cause secondary changes that are outcome rather than the cause of the behavioral changes. Therefore, we subjected the hippocampus from 6-month-old WT and APP^NL-G-F^ mice to nuclei isolation and capture after exercise training for 3 months. The snRNA-seq and snATAC-seq analyses were performed in different batches with different mice. For each analysis, hippocampi from 3 mice were pooled together before nuclei isolation and capture. After quality control, we obtained 31,793 nuclei for snATAC-seq and 34,950 nuclei for snRNA-seq, which were clustered into 6 major cell types, including excitatory neurons (EN), inhibitory neurons (IN), oligodendrocytes (OLG), oligodendrocyte progenitor cells (OPC), astrocytes (AST), and microglia (MG) (**Fig 1j-k**). Exercise did not cause obvious changes in the overall cellular composition in WT or APP^NL-G-F^ mice (**Suppl Fig S2a**). We focus on the exercise effect (APP_EX vs. APP_RT and WT_EX vs. WT_RT comparisons) and the amyloid effect (APP/RT vs. WT/RT comparison). Differentially expressed genes (DEGs) from these comparisons were identified from each cell type. Exercise led to over 3 - 6 folds more DEGs in APP mice than in WT mice in most cell types (**Suppl Fig S2b, Suppl Table S1-6**), consistent with more robust improvement of the cognitive functions in APP^NL-G-F^ mice compared to WT mice.

Exercise caused a predominant upregulation of gene expression, while amyloid led to a predominant downregulation of gene expression (**Suppl Fig S2b-c**). DEGs altered by exercise or amyloid were enriched in different functional pathways in a cell type-specific manner (**Suppl Fig S2c-d, Suppl Table S7-9**). For example, DEGs in microglia were enriched in the complement and IL5/IL6 signaling pathways; those in oligodendrocytes were enriched in prostaglandin signaling; those in OPCs were enriched in the IL12/STAT4 signaling; those in astrocytes were enriched in lipid metabolism, angiogenesis, and cytoskeletal regulation; and those in neurons were enriched calcium signaling, protein translation, and mRNA processing (**Suppl Fig S2d**). Notably, the receptor tyrosine kinase signaling, especially insulin, c-KIT, epidermal growth factor receptor (EGF), and the downstream phosphoinositide 3-kinases (PI3K) signaling pathways were universally enriched across multiple cell types (**Suppl Fig S2d**).

### Exercise counteracts amyloid-dependent transcriptomic changes in growth factor signaling

The exercise-induced genes (APP_EX vs. APP_RT) showed a robust negative correlation with the amyloid-induced genes (APP_RT vs. WT_RT) across all cell types (**Fig 2a**). Over 833 reversed DEGs were found in at least one cell type, with significant changes in both comparisons but in opposite directions. Most of these DEGs were upregulated by exercise and downregulated by amyloid (**Fig 2b**), suggesting that exercise ‘reversed’ the amyloid-induced transcriptomic changes by activating transcription. Interestingly, less than 9 reversed genes were shared by any 4 clusters (**Fig 2c**), suggesting that the reversal effects are highly cell type-specific at the gene level. However, at the pathway level, the EGF receptor (EGFR) and insulin pathways stood out as the top common functional pathways with the reversal pattern across most cell types (**Fig 2d-f, Suppl Table S10-11**).

**Figure 2.**
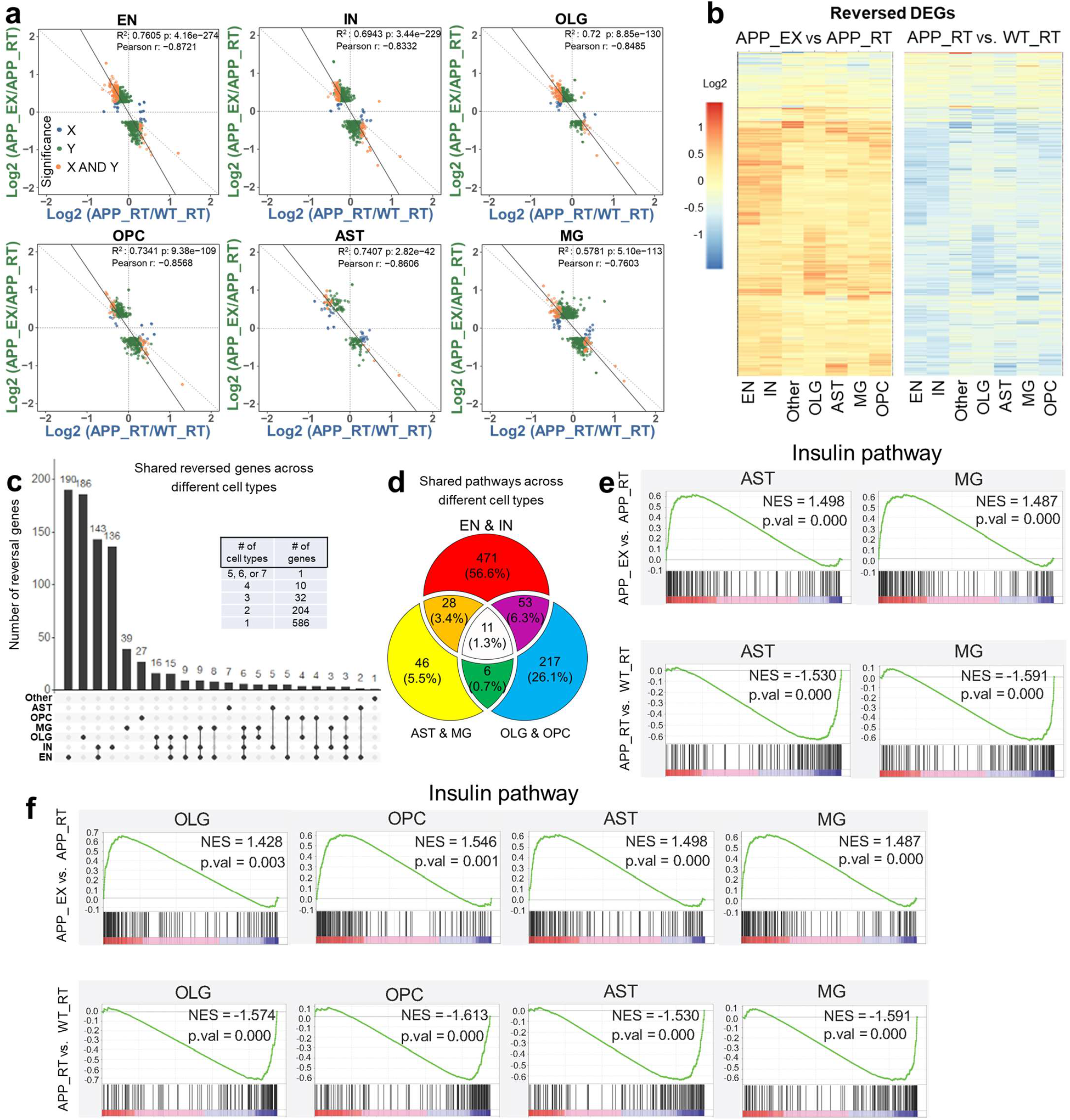
Exercise counteracts amyloid-dependent transcriptomic changes. (**a**) Scatter plot of gene expression showing a negative correlation between exercise effects (APP_EX vs. APP_RT) and amyloid effects (APP_RT vs. WT_RT) across different cell types. (**b**) Heat map of 833 differentially expressed genes (DEGs) with the reversed pattern in at least one cell type. (**c**) Number of overlapping reversed DEGs across different cell types. (**d**) Overlapping pathways enriched in reversed DEGs across different cell types. (**e-f**) GSEA analysis of the insulin signaling pathway, a top common enriched pathway in reversed genes in different cell types.

The EGFR/insulin pathway was suppressed by amyloid and upregulated by exercise (**Fig 3a** and **Suppl Fig S3**). Stxbp1 is a top gene in neurons with a reversed expression pattern within the EGFR/insulin pathway and encodes a syntaxin-binding protein involved in synaptic vesicle cycling. RNAscope verified that hippocampal Stxbp1 was suppressed by amyloid and upregulated by exercise (**Fig 3b-c**). EGFR and insulin signaling share many downstream players, including PI3K, AKT, and MAPK (**Fig 3d-e**). The EGFR/insulin signaling pathway functions downstream of several growth factors, such as insulin-like growth factor (IGF), fibroblast growth factors (FGF), hepatocyte growth factor (HGF), transforming growth factor (TGF), vascular endothelial growth factor (VEGF), and platelet-derived growth factor (PDGF) (**Fig 3e**). In summary, exercise counteracts amyloid-induced repression of the growth factor signaling in multiple cell types.

**Figure 3.**
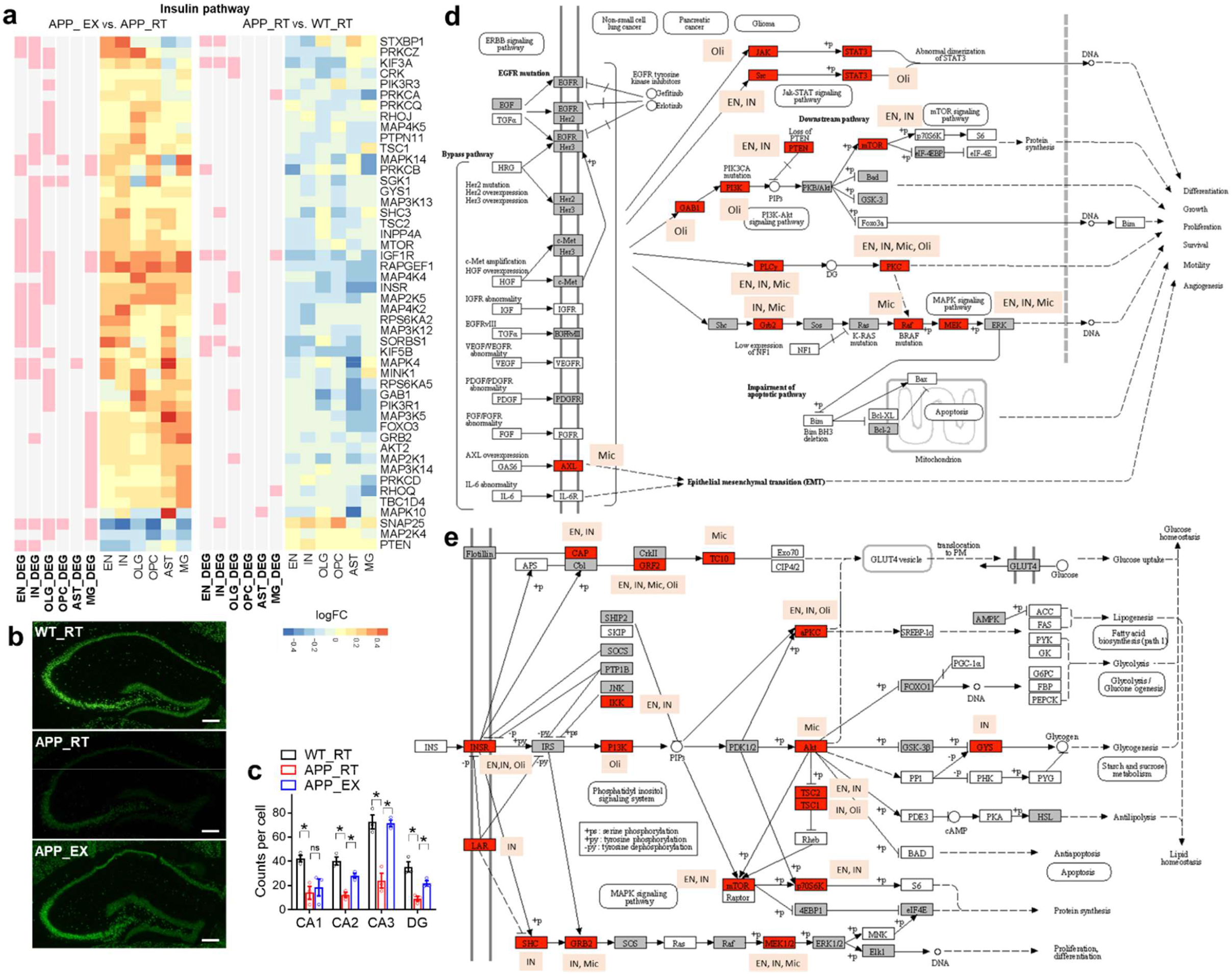
Exercise activates the EGFR/insulin signaling. (**a**) Heat map of reversed DEGs within the insulin signaling in different cell clusters. (**b-c**) RNAscope analysis of Stxbp1, a gene of the insulin/EGFR pathway with known function in neurotransmission and a robust reversed expression pattern in EN. Scale bar, 200 µm. n = 3 mice. Bar graphs show the mean with S.E.M. Asterisks indicate significant differences by t-test. ns, non-significant. (**d**) Some of the DEGs in the EGFR pathway. (**e**) Some of the DEGs in the insulin pathway. Images were generated from the KEGG pathway database.

### Exercise counteracts amyloid-induced transcriptional regulatory networks

snATAC-seq identified differentially accessible regions (DARs) in response to exercise or amyloid deposition in each cell type (**Suppl Fig S4a**). The top enriched pathways and motifs in these DARs were cell type-specific (**Suppl Fig S4b-c**). Among the enriched transcription factors (TFs) are those related to growth factor signaling, cell proliferation, and neuron differentiation, including EGR1, MYB, ATOH1, and ASCL1 (**Suppl Fig S4c**). Consistent with transcriptomics data, exercise-induced and amyloid-induced genome accessibility changes displayed negative correlations across all cell types (**Fig S4d**). Unlike the transcriptomics data, DARs with the reversal phenotype were more evenly distributed in both directions (**Fig 4a**). However, motif analyses of these reversed DARs revealed direction-specific transcription regulator networks centered on TFs (**Fig 4b**). DARs upregulated by amyloid and downregulated by exercise were referred to as “U>D” (from ‘Upregulation’ to ‘Downregulation’), while those DARs downregulated by amyloid and upregulated by exercise were referred to as “D>U” (from ‘Downregulation’ to ‘Upregulation’). TFs involved in growth and differentiation, such as EGR1, MYB, MEF2, and ASCL1, show direction-specific enrichment in neurons (**Fig 4b**). These TFs are downstream of growth factors, PI3K, or MAPK signaling pathways and are involved in synaptic plasticity or cell growth^19–22^. WT1 and NEUROG2, transcription factors in cell growth and neurogenesis^23–25^, were enriched in the D>U DARs in both excitatory neurons and inhibitory neurons. ATOH1, the transcription factor essential for cerebellar granule cell formation^26^, was enriched in D>U DARs of excitatory neurons, and U>D DARs of inhibitory neurons (**Fig 4b**). DARs of both directions were enriched with the ETS family of TFs in microglia, the SOX family of TFs in oligodendrocytes, and the LHX family of TFs in astrocytes (**Fig 4b**).

**Figure 4.**
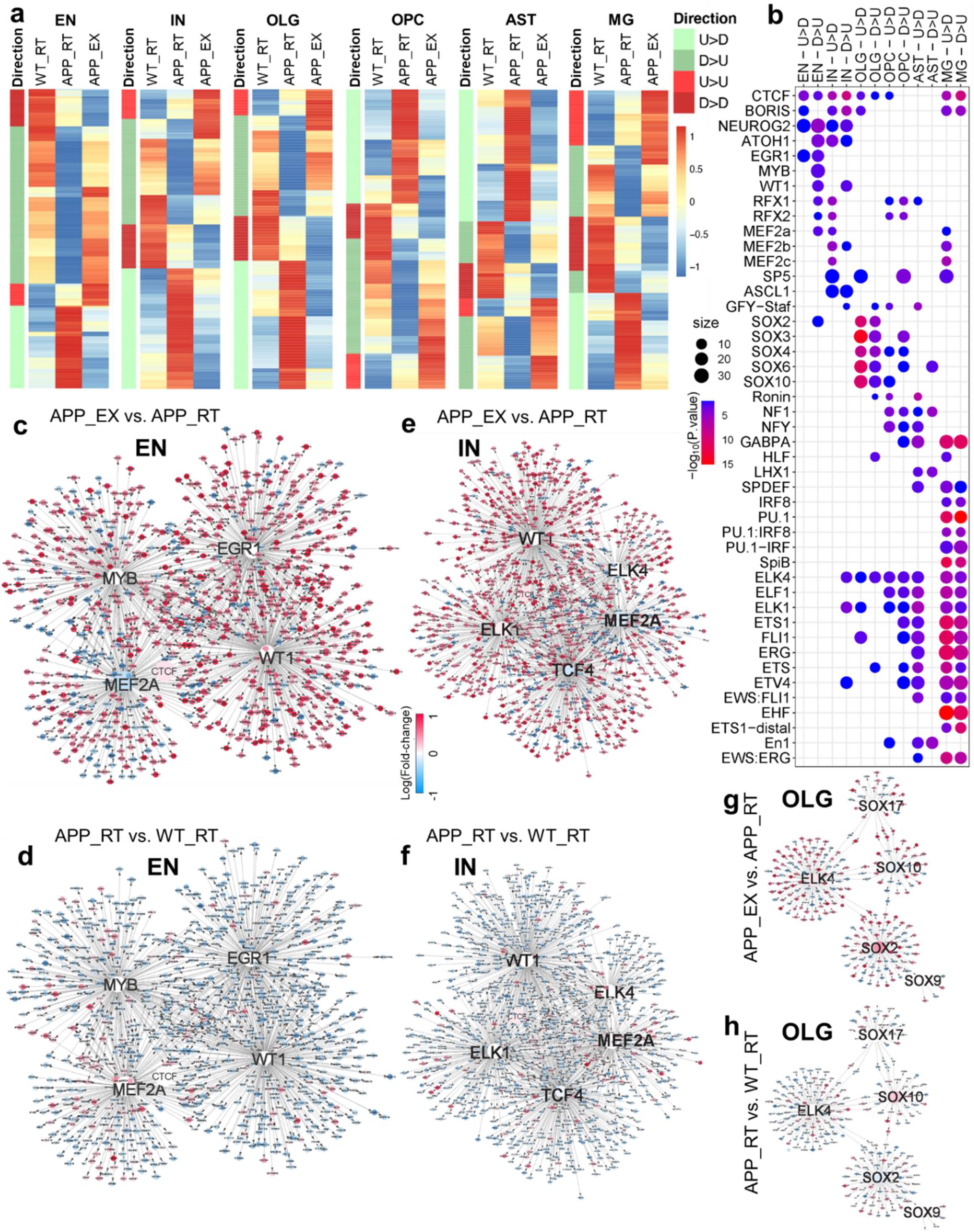
Exercise stimulates cell type-specific transcriptional regulatory networks. (**a**) Heat map of the relative levels of the reversed differentially accessible regions (DARs) in each cluster. DARs upregulated by APP^NL-G-F^ (APP_RT vs. WT_RT) and downregulated by exercise (APP_EX vs. APP_RT) were referred to as “U>D”, while those DARs downregulated by APP^NL-G-F^ and upregulated by exercise were referred to as “D>U”. (**b**) Top enriched motifs in the DARs in each cell cluster. (**c-h**) Top network showing direction-specific enrichment in EN, IN, and OLG populations.

Transcriptional regulatory network analysis of the excitatory neurons suggested that EGR1, WT1, MYB, and MEF2 are central TFs that reversed amyloid-mediated transcriptomic changes by activating transcription in response to exercise (**Fig 4c-d** and **Suppl Fig S5**). The downstream genes of these central TFs in the network overlapped significantly with genes in the EGFR/insulin pathway (**Suppl Table S12-13**). Many transcription factors can serve as transcription activators and repressors in a context-dependent manner. Therefore, it makes sense that the same transcription factor network can drive opposite reversal directions. In inhibitory neurons, ELK1/4 and TCF4 replace EGR1/MYB as key TFs working with MEF2A and WT1 for exercise-induced transcriptional remodeling (**Fig 4e-f** and **Suppl Fig S6**). By comparison, the network analysis suggests that the SOX family TFs work with ELK4 to drive the reversal of the APP-induced oligodendrocyte over-maturation by exercise (**Fig 4g-h** and **Suppl Fig S7**), in line with the known role of the SOX family in oligodendrocyte differentiation^27^. Similar patterns were observed in other cell types, with cell type-specific TFs driving distinct downstream genes that converge on the EGFR/insulin pathway (**Suppl Fig S8-S10** and **Suppl Table S12-13**). In summary, exercise stimulates cell type-specific transcriptional regulatory networks, counteracting amyloid-induced transcriptomic changes by activating growth factor signaling.

One cellular manifestation of growth factor signaling activation is neurogenesis. Granule cells (GC) in the dentate gyrus (DG) of the hippocampus constitute the primary niche for adult hippocampal neurogenesis^28^. A more focused analysis of the snRNA-seq data within the hippocampal excitatory neurons revealed that exercise causes a more drastic gene expression change in the GC population than the pyramidal cell population (**Fig 5a-c**), leading to a higher proportion of granule cells within the EN cluster (**Fig 5d**). Trajectory analysis of neurons did not recapitulate the neurogenesis process, probably due to the scarcity of nascent neurons in the adult brain. Therefore, we sought to resolve the snRNA-seq data spatially to address whether exercise-induced differences show spatial preference towards DG GCs. We integrated the EN snRNA-seq data with the previous spatial transcriptomic data^29^ to group EN nuclei into sub-hippocampal regions, such as DG (8813 nuclei) and CA1 (3379 nuclei) (**Suppl Fig S11**). We found that exercise increased the GC proportion in the DG, but not IN proportion in the DG (**Fig 5e**). Production of immature GCs (imGCs) is a hallmark of adult hippocampal neurogenesis. To predict imGCs, we applied the logistic regression model trained on mice prototype imGCs (with a gene signature of Ascl1, Dcx, Tubb3, Neurod1, and Tbr1)^30^ to our snRNA-seq data. The imGCs proportion was reduced by amyloid, which was rescued by exercise (**Fig 5f**). These results suggest that multiple growth factor signaling and neurotrophic pathways participate in exercise-stimulated neurogenesis in the DG.

**Figure 5.**
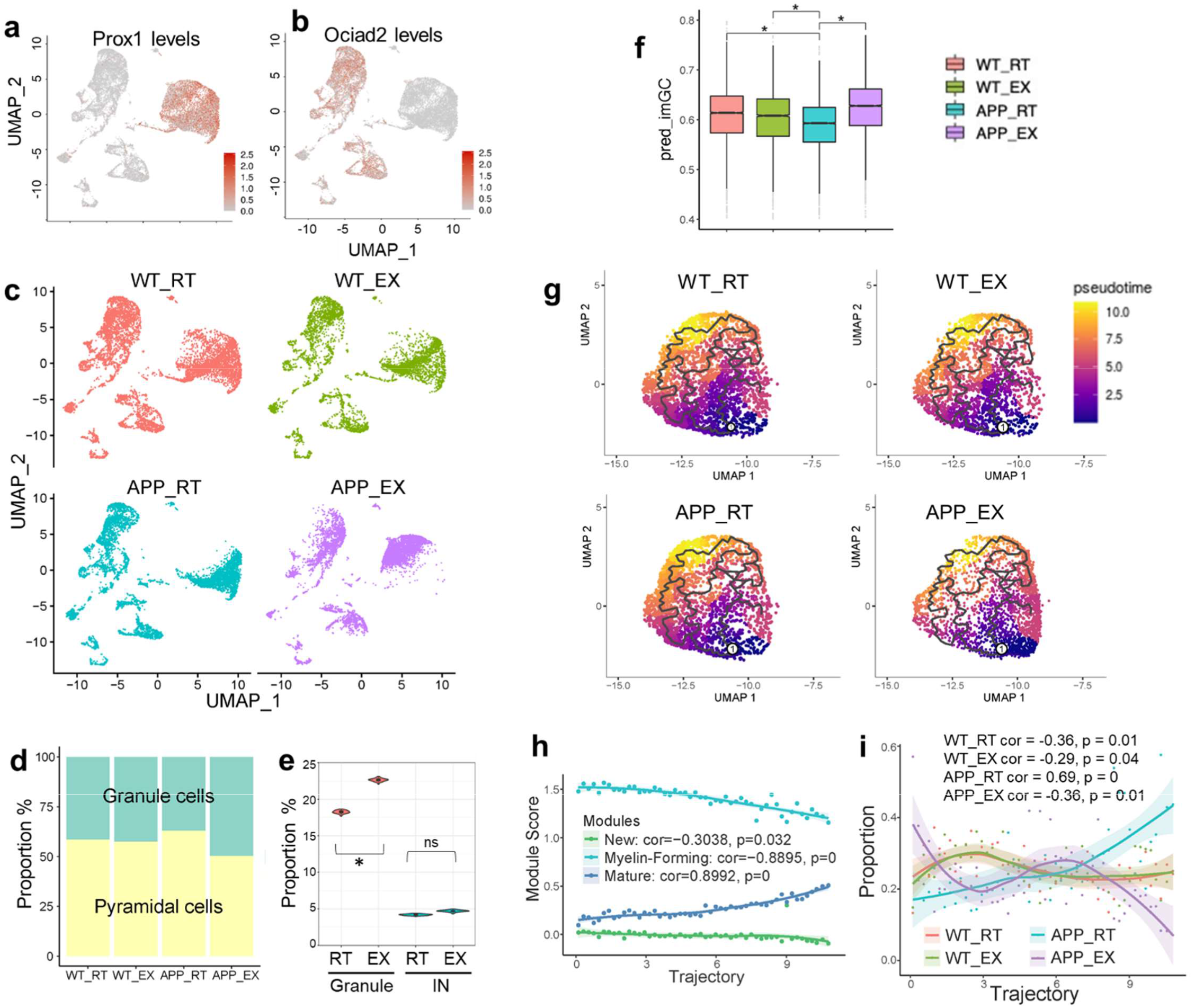
Exercise impacts excitatory neurons and oligodendrocytes. (**a-b**) UMAP of snRNA-seq data in EN sub-clusters: Prox1+ granule cells and Ociad2+ pyramidal cells. Expression is based on library-size-normalized log values. (**c**) UMAP of hippocampal EN sub-clusters in each individual group. (**d**) The proportion of granule and pyramidal cells within the EN cluster in each group. (**e**) Proportions of granule cells and INs in the DG. (**f**) Box plot of predicted GC immature score (imGC). 514 of 534 genes in the model were found in our data, and missing genes had only small weights in the model. The log-transformed and max-normalized counts matrix were taken as the input to predict the final imGC score from the logistic regression model. Center line, median; box limits, upper and lower quartiles; whiskers, 1.5x interquartile range; points, outliers. Asterisks indicate significant differences. ns, non-significant. (**g**) UMAP of the OLG for trajectory analysis. (**h**) Module score along the pseudotime trajectory of the OLG. (**i**) Distribution of each group on the OLG trajectory.

In addition to neurons, oligodendrocytes stood out as another cell type with a growth and proliferation phenotype amenable to exercise. We integrated snRNA-seq and snATAC-seq and took the co-embedding space for trajectory analysis^31^ (**Fig 5g**). The trajectory recapitulated oligodendrocyte maturation because the expression signature of the new, myelin-forming, and mature oligodendrocyte showed a monotonic correlation with the pseudo-time scale (**Fig 5h**). Amyloid increased the proportion of mature oligodendrocytes (**Fig 5i**), which is in line with previous reports that amyloid oligomers promote oligodendrocyte differentiation and lead to thicker myelin^32,33^. We speculate that this over-maturation phenotype could be due to senescence or a lack of replenishment. Interestingly, exercise reduced the mature oligodendrocyte proportion while increasing myelin-forming and new oligodendrocyte proportions (**Fig 5i**).

### Growth factor signaling contributes to the memory-enhancing effects of exercise

The single-nucleus multi-omics suggest that several TFs-centered networks contribute to exercise-stimulated activation of growth factor signaling pathways in different cell types. To address whether the EGFR-related growth factor signaling is required for exercise-mediated cognitive improvement, we administered EGFR inhibitor Gefitinib at 50 mg/kg and PI3K inhibitor Wortmannin at 0.5 mg/kg through oral gavage once every other day from 4 to 8 months old in APP^NL-G-F^mice while they were subjected to wheel-running starting at 4 months old. These pharmacologic manipulations did not affect the wheel-running exercise volume (**Fig 6a-b**) or body weight (**Fig 6c**). However, the drugs efficiently blunted the exercise-mediated cognitive improvement in the object-in-place test (**Fig 6d-e**), Y-maze test (**Fig 6f-g**), social memory (**Fig 6h-j**), and Morris water maze test (**Fig 6k-l**) without affecting sociability (**Fig 6i**) or the total locomotor activity during these tests. The inhibitors did not alter anxiety-related behaviors in the elevated plus maze test, open field arena test, or light-dark test (**Fig 6m-o**). Interestingly, the inhibitors also blunted exercise-induced improvement in amyloid pathology (**Fig 6p-q**), suggesting that EGFR signaling activation and its related anabolic stimulation are required for exercise-induced amyloid clearance and cognitive benefits.

**Figure 6.**
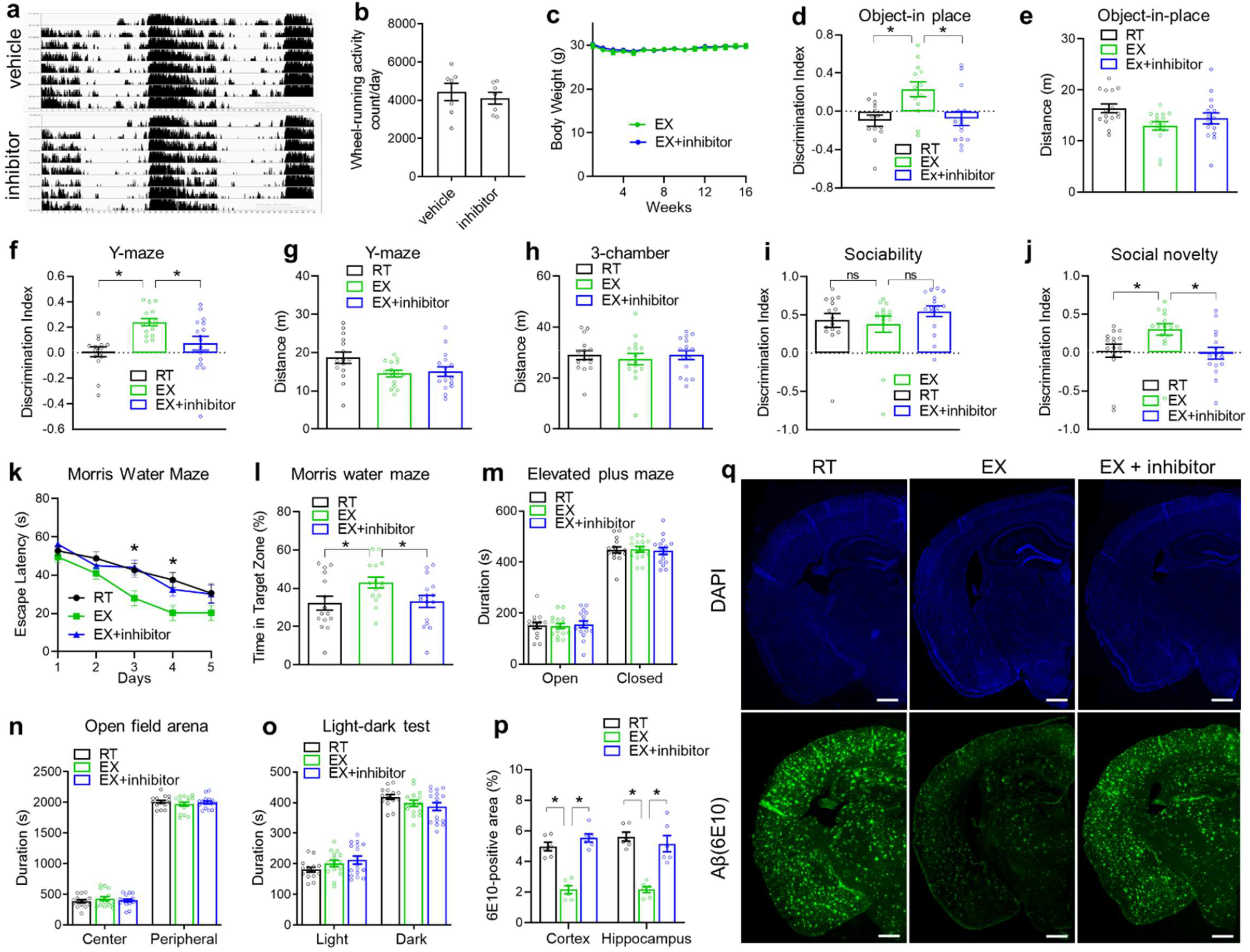
EGFR and PI3K signaling are required for the cognitive-improving effects of exercise. (**a**) Average actogram profiles of wheel-running activity of 8-month-old male mice treated with EGFR inhibitor Gefitinib and PI3K inhibitor Wortmannin through oral gavage once every other day from 4 to 8 months old. Mice were simultaneously subjected to wheel-running from 4 to 8 months old. (**b**) Average daily wheel-running activity (n = 7 cages per group with 2 mice per cage). (**c**) Body weight. (**d-o**) Object-in-place, Y-maze, 3-chamber sociability and social memory, Morris water maze, elevated plus maze, open field arena, and light-dark tests (n = 15 mice for RT, 15 mice for EX, and 16 mice for EX + inhibitor). 2 RT, 2 EX and 1 EX+inhibitor mice were excluding due to discrimination index were greater or smaller than ± 0.7 (**p-q**) Immunostaining of β-amyloid in mice treated with inhibitors. Scale bar: 600 µm. n = 6 mice per group. All bar graphs show the mean with S.E.M. Asterisks indicate significant differences by one-way ANOVA with Fisher’s LSD multiple comparisons except Morris water maze was analyzed with repeated-measure 2-way ANOVA with Fisher’s LSD multiple comparisons where asterisks indicate differences between RT vs. EX and EX + inhibitor vs. EX.

We sought to identify the upstream signals for exercise-induced growth factor signaling. Among ligands for the EGFR family^34^, heparin-binding EGF-like growth factor (HB-EGF) stood out with significant gene expression upregulation in mouse muscles^35^ and human muscles after physical exercise^36^ (**Fig 7a**). HB-EGF gene expression is also upregulated in blood cells after exercise in human blood^37^. In a proteomics analysis of human blood samples, HB-EGF is the only detectable EGFR ligand upregulated by exercise^38^ (**Fig 7b**). HB-EGF is known to cross the blood-brain barrier^39^. We confirmed that the blood HB-EGF levels were elevated after chronic exercise in APP^NL-G-F^ mice (**Fig 7c**). To address whether HB-EGF has cognitive benefits, we intranasally administered HB-EGF in sedentary APP^NL-G-F^ mice at 3 ug/mouse once every other day from 4 to 7 months old. HB-EGF did not affect body weight (**Fig 7d**) but improved cognitive functions in the object-in-place test (**Fig 7e-f**), Y-maze test (**Fig 7g-f**), and social memory (**Fig 7i**) without affecting the sociability (**Fig 7j**) or locomotor activity during the three-chamber test (**Fig 7k**). HB-EGF also reduced escape latency during the multiple-day Morris water maze test (**Fig 7l**) but did not cause significant differences in the probe test on the last day (**Fig 7m**). HB-EGF did not alter anxiety-related behaviors in the open field arena test or light-dark test (**Fig 7n-o**) but reduced the overall beta-amyloid deposition (**Fig 7p-q**). These results suggest chronic intranasal HB-EGF treatment in mice can ameliorate amyloid-induced cognitive decline and reduce amyloid deposition.

**Figure 7.**
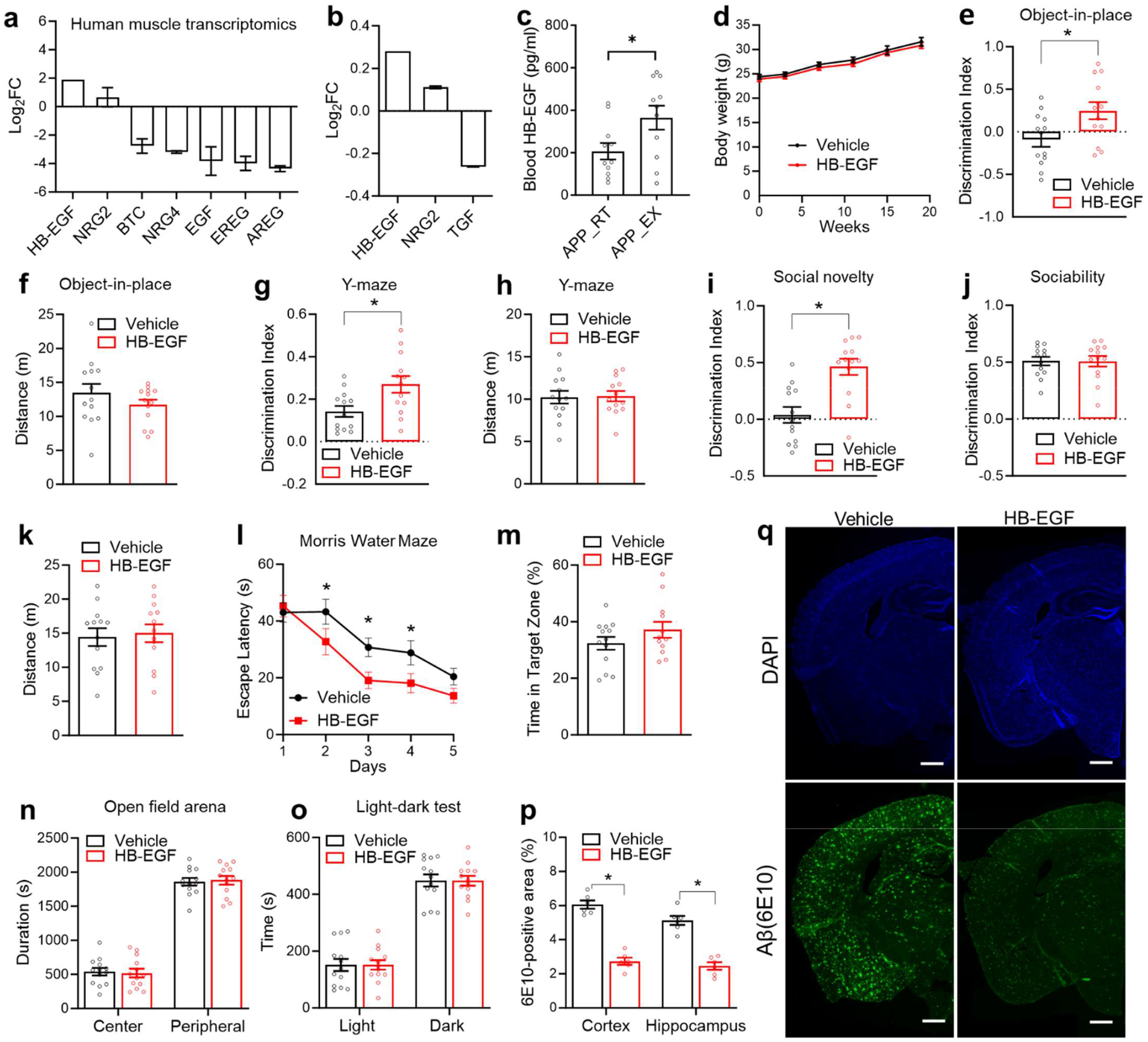
Intranasal HB-EGF mimics exercise-induced cognitive improvement. (**a**) Replot of gene expression levels of EGF family members in human skeletal muscles after long-term exercise training from a published transcriptomic dataset^36^. (**b**) Replot of protein levels of EGF family members in the human blood after long-term exercise training from a published proteomics dataset^38^. (**c**) Serum HB-EGF levels in APP^NL-G-F^ male mice after chronic wheel-running exercise for 6 months (n = 11 mice for RT, and 11 mice for EX). (**d**) Body weight gain during intranasal HB-EGF administration in female APP^NL-G-F^ mice. Administration started at 4 months old, with once every other day (n = 13 mice for the vehicle; 13 mice for HB-EGF). (**e-o**) Object-in-place, Y-maze, 3-chamber sociability and social memory, Morris water maze test, open field arena test, and light-dark test in female mice at 7-8-months old after chronic HB-EGF administration (n = 13 mice for the vehicle; 13 mice for HB-EGF; one mouse from the HB-EGF group was excluding from MWM due to low mobility). (**p-q**) Immunostaining of β-amyloid in 8-month-old female mice treated with intranasal HB-EGF for 4 months. Scale bar: 600 µm. n = 6 mice per group. All bar graphs show the mean with S.E.M. Asterisks indicate significant differences by one-way ANOVA with Fisher’s LSD multiple comparisons or repeated-measure ANOVA with Fisher’s LSD multiple comparisons.

## DISCUSSION

Our results offer a comprehensive overview of transcriptomic and chromatin accessibility changes across different cell types within the mouse hippocampus in response to chronic voluntary exercise. Two recent publications present snRNA-seq analyses conducted on mouse brains. One study examined the whole brain following 12 months of voluntary wheel-running exercise^40^, while the other focused on the hippocampus after 4 weeks of wheel-running exercise^41^. These investigations were conducted on wild-type mice without amyloid deposition and did not include snATAC-seq analyses. Our utilization of APP^NL-G-F^ mice, coupled with snATAC-seq integration, illuminates the upstream transcriptional factor networks governing hippocampal responses to exercise in the presence of amyloid deposition. Our profiling reveals that exercise reverses amyloid-induced transcriptomic alterations by activating gene transcription. Exercise-induced transcriptional regulatory networks show specificity to cell types for upstream transcription factors. Yet the downstream target genes collectively converge on growth factor signaling pathways, particularly the EGFR/insulin pathway, which is associated with elevated HB-EGF levels in the blood. The cognitive benefits of exercise are blocked by pharmacological inhibition of EGFR/insulin signaling, while chronic intranasal administration of HB-EGF enhances memory function in sedentary APP^NL-G-F^ mice. Therefore, the insights gained from single nucleus multi-omics analysis of exercise effects on the brain have opened the door to a potential therapeutic approach for AD by activating growth factor signaling.

Growth factors, including BDNF, IGF-1, VEGF, and GH, have been implicated in the neurotrophic or synaptogenic effects of exercise^42,43^. Our findings suggest that HB-EGF is a novel growth factor involved in the process. Our results align with prior studies showing HB-EGF administration can enhance the generation of new neurons or oligodendrocytes^44,45^. Consistently, HB-EGF was shown to interact with APP and promote cellular neuritogenesis^46^. These findings do not rule out other signaling pathways in exercise-induced cognitive improvement. Interestingly, EGFR inhibitors were reported to have beneficial effects in AD, with some conflicting results ^47,48^. We find that a combined EGFR inhibitor and PI3K inhibitor blocked the effects of exercise training, which may act independently of the baseline effects of the EGFR inhibitor itself. EGFR effects on AD appear to be age-dependent and mediated by glial cells^47^. There is currently no available data on the effects of EGFR inhibitors in the APP^NL-G-F^ mouse model at the baseline. Although we focus on HB-EGF based on muscle and blood omics datasets available in the literature, many brain cell types can produce EGF factors, which could be an additional source of elevated EGFR signaling. Further research is needed to elucidate the source of HB-EGF and the effects of EGFR inhibitors or agonists on AD progression.

The current study has several limitations. Firstly, the pharmacokinetics and pharmacodynamics of intranasal HB-EGF administration remain unclear. We speculate that intranasal administration may result in brain-enriched distribution, activating EGFR signaling and potentially mimicking the effects of exercise training in promoting neuritogenesis or neurogenesis. Secondly, it is uncertain whether treatment with Gefitinib and Wortmannin reduces EGFR and PI3K signaling in the hippocampus and whether this would negatively impact cognitive functions or amyloid pathology in the baseline condition without exercise training. Hence, it cannot be conclusively stated that the effects of EGFR/PI3K inhibition are attributed explicitly to exercise. Lastly, hippocampal samples from three mice were pooled for the single-nuclei omics analysis to enhance cost efficiency, albeit at the expense of statistical power. Future advancements in techniques may enable more cost-efficient comprehensive profiling. Despite these limitations, the identification of EGFR signaling from non-biased omics datasets, the EGFR/PI3K inhibitors-mediated abrogation, and the intranasal HB-EGF-mediated recapitulation of exercise-induced cognitive improvements and amyloid pathology collectively support a positive role of the EGFR signaling pathway in the cognitive benefits of exercise in the presence amyloid deposition.

A fundamental function of growth factor signaling is stimulating anabolic metabolism^49^, which may or may not lead to cellular proliferation or organellar growth. Our results suggest that anabolic resistance might be a prevalent feature of the aging brain, contributing to cognitive decline and the pathogenesis of AD, but potentially mitigated by exercise. Thus, the opposing dynamics of growth and senescence could explain the inverse correlation between cancer and AD observed in the elderly human population ^50^.

## ACKNOWLEDGEMENT

We thank Drs. Hesong Liu, Gabriella Perez, and Joanna Jankowsky at Baylor College of Medicine (BCM) for helpful discussions, and Fei Peng at BCM for technical assistance in drug administration. We thank Drs. Yumei Liu and Rui Chen at the BCM Single Cell Genomics Core (supported by NIH S10OD025240 and CPRIT RP200504) for single nuclei capture and Dr. Daniel Kraushaar at the BCM Genomic and RNA Profiling Core (supported by NIH NCI P30CA125123 and CPRIT RP200504) for sequencing. We thank Dr. Sonia Villapol and Morgan Holcomb at Houston Methodist Hospital for their effort in histology analysis. The authors’ laboratories are supported by NIH AG069966, AG070687, and DK111436. The authors are grateful to the Alzheimer’s Association (AARG-21-847542), John S. Dunn Foundation, Texas Medical Center Digestive Diseases Center (P30DK056338), the SPORE program in lymphoma (P50CA126752), and the Gulf Coast Center for Precision Environmental Health (GC-CPEH, P30ES030285).

## AUTHOR CONTRIBUTIONS

XL performed most behavioral assays and amyloid histology analysis; CL performed integrated snRNA-seq/snATAC-seq analysis; WL executed the exercise protocol, isolated nuclei, and performed RNAscope analysis; YD performed the initial snRNA-seq analysis; CG performed the spatial transcriptomics-related analysis; WZ and JL performed some of the behavioral assays; VCC assisted in preparing figures and language editing; HZ provided and advised on the mouse model; UK, DG, ZH, HC assisted or advised on data analyses or data interpretation. ZL guided bioinformatics analysis; YW supervised bioinformatics analysis; ZS and ZL conceived the study and obtained funding; YW and ZS wrote the manuscript with input from other authors.

## CONSENT STATEMENT

No human subjects were involved. Consent was not necessary. The publication of the described work is approved by all authors.

