## Supplemental Information for "Multi-omics delineate growth factor network underlying exercise effects in an Alzheimer’s mouse model"

#### SUPPLEMENTARY TABLES

**Supplemental Table S1. RNA in excitatory neurons**
**Supplemental Table S2. RNA in inhibitory neurons.**
**Supplemental Table S3. RNA in oligodendrocytes.**
**Supplemental Table S4. RNA in oligodendrocyte progenitor cells.**
**Supplemental Table S5. RNA in astrocytes.**
**Supplemental Table S6. RNA in microglia.**
**Supplemental Table S7. GSEA results of DEGs for APP_RT vs. WT_RT**
**Supplemental Table S8. GSEA results of DEGs for APP_EX vs. APP_RT**
**Supplemental Table S9. GSEA results of DEGs for DEGs WT_EX vs. WT_RT**
**Supplemental Table S10. GSEA results of the EGFR signaling pathway.**
**Supplemental Table S11. GSEA results of the insulin signaling pathway.**
**Supplemental Table S12. Overlap of EGFR signaling with TFs targets genes in network analysis.**
**Supplemental Table S13. Overlap of Insulin signaling with TFs targets genes in network analysis.**

#### SUPPLEMENTARY FIGURE LEGENDS

**Supplemental Figure S1. Behavioral analysis of WT and APP$^{NL\text{-}G\text{-}F}$ mice.**
(**a**) Average daily exercise volume per mouse. n = 8 WT mice, n = 9 APP$^{NL\text{-}G\text{-}F}$ mice. (**b**) Body weight, n = 12 mice for WT_RT, n = 8 mice for WT_EX; 10 mice for APP_RT; 9 mice for APP_EX. (**c**) Total distance in the standard 2-object OLT (n = 12 for WT_RT; 8 for WT_EX; 10 for APP_RT; and 9 for APP_EX). (**d**) Total distance in the Y-maze test (n = 12 for WT_RT; 8 for WT_EX; 10 for APP_RT; and 9 for APP_EX). Bar graphs show the mean with S.E.M. Asterisks indicate significant difference by two-way ANOVA. ns, not significant.

**Supplemental Figure S2. Exercise activates growth factor signaling.** (**a**) Proportions of major cell clusters based on snRNA-seq and snATAC-seq. (**b**) The number of DEGs from the indicated comparisons. (**c**) The number of significantly enriched pathways in DEGs from the indicated comparisons. (**d**) List of significantly enriched pathways in DEGs from the indicated comparisons.

**Supplemental Figure S3. DEGs with the reversed pattern from snRNA-seq analysis.**
(**a**) GSEA analyses of the EGFR signaling pathway, a top common enriched pathway in reversed DEGs in different cell types. (**b**) Heat map of reversed DEGs within the EGFR signaling in different cell types.

**Supplemental Figure S4. DARs from snATAC-seq analysis.**
(**a**) The number of DARs from the indicated comparisons. (**b**) Number of significantly enriched pathways in DARs from the indicated comparisons. (**c**) List of significantly enriched motifs in the indicated DARs. (**d**) Scatter plot of chromatin regions with negative correlations between the exercise-induced (APP_EX vs. APP_RT) and amyloid-induced (APP_RT vs. WT_RT) accessibility changes across different cell clusters.

**Supplemental Figure S5. Transcriptional regulatory network in excitatory neurons.**
**Supplemental Figure S6. Transcriptional regulatory network in inhibitory neurons.**
**Supplemental Figure S7. Transcriptional regulatory network in oligodendrocytes.**
**Supplemental Figure S8. Transcriptional regulatory network in oligodendrocyte precursor cells.**
**Supplemental Figure S9. Transcriptional regulatory network in astrocytes.**
**Supplemental Figure S10. Transcriptional regulatory network in microglia.**

**Supplemental Figure S11. Spatial expression reconstruction.**
Heat map of DG and CA1/2 landmark genes, Prox1 and Ociad2, in WT_RT mouse hippocampus after spatial expression reconstruction according to a previous spatial transcriptomics dataset.

### Supplementary Figure S1

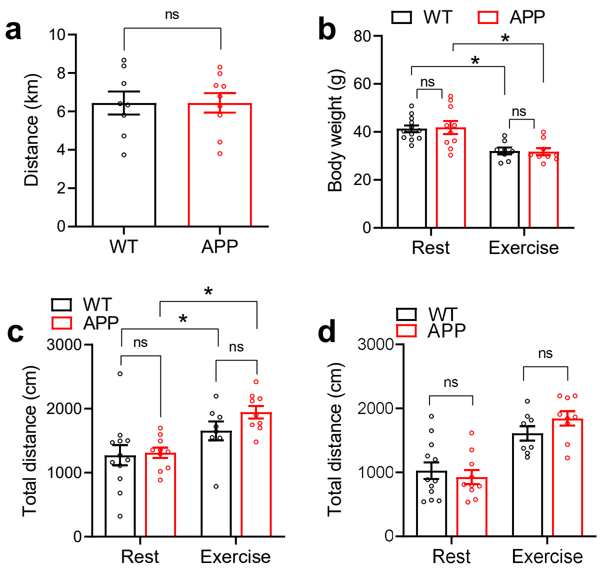

### Figure S2

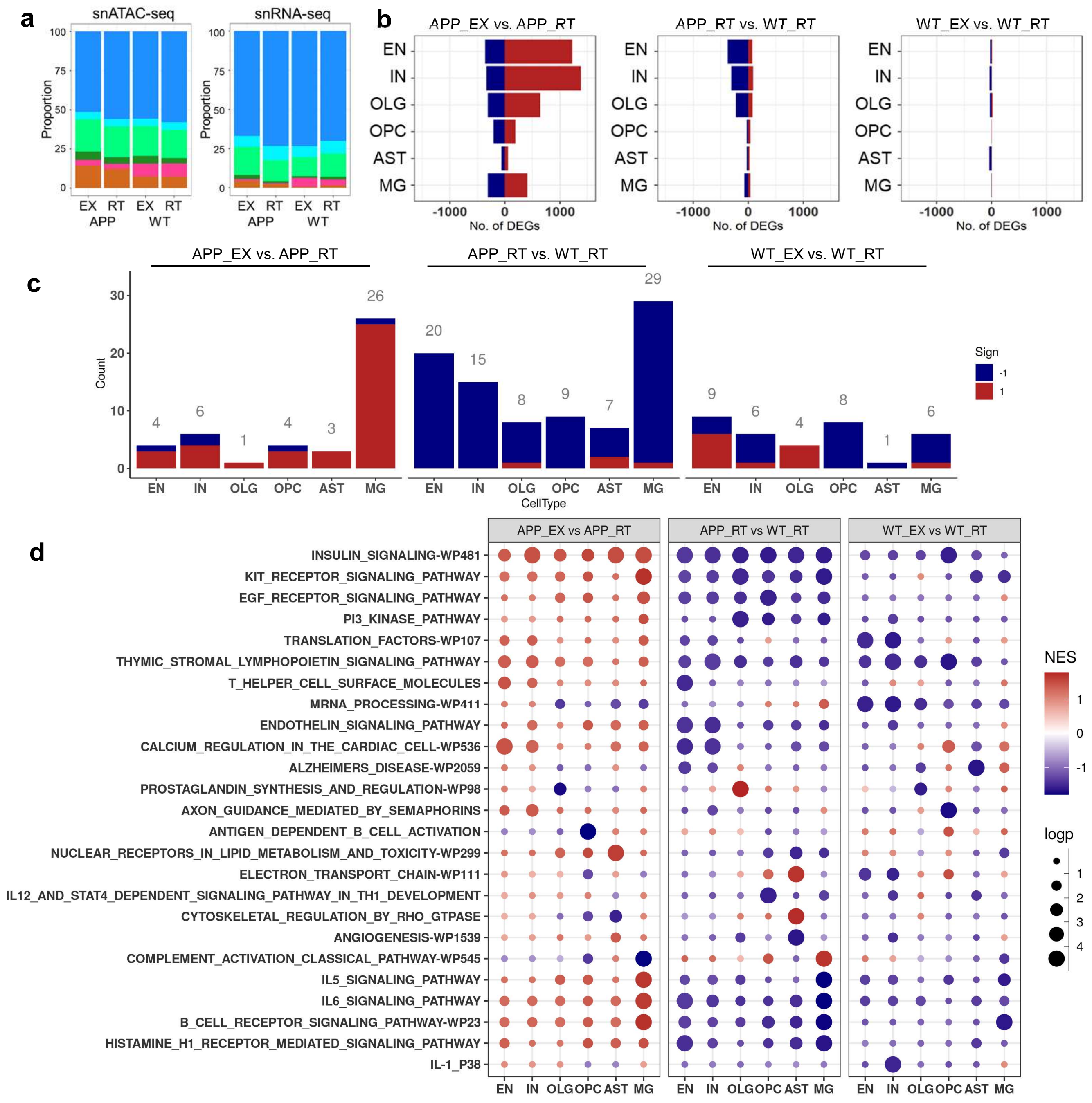
a
snATAC-seq
snRNA-seq
Proportion
100
75
50
25
0
EX RT EX RT
APP WT
b
APP_EX vs. APP_RT
APP_RT vs. WT_RT
WT_EX vs. WT_RT
EN
IN
OLG
OPC
AST
MG
-1000
0
1000
No. of DEGs
c
APP_EX vs. APP_RT
APP_RT vs. WT_RT
WT_EX vs. WT_RT
Count
CellType
Sign
-1
1
d
APP_EX vs APP_RT
APP_RT vs WT_RT
WT_EX vs WT_RT
INSULIN_SIGNALING-WP481
KIT_RECEPTOR_SIGNALING_PATHWAY
EGF_RECEPTOR_SIGNALING_PATHWAY
PI3_KINASE_PATHWAY
TRANSLATION_FACTORS-WP107
THYMIC_STROMAL_LYMPHOPOIETIN_SIGNALING_PATHWAY
T_HELPER_CELL_SURFACE_MOLECULES
MRNA_PROCESSING-WP411
ENDOTHELIN_SIGNALING_PATHWAY
CALCIUM_REGULATION_IN_THE_CARDIAC_CELL-WP536
ALZHEIMERS_DISEASE-WP2059
PROSTAGLANDIN_SYNTHESIS_AND_REGULATION-WP98
AXON_GUIDANCE_MEDIATED_BY_SEMAPHORINS
ANTIGEN_DEPENDENT_B_CELL_ACTIVATION
NUCLEAR_RECEPTORS_IN_LIPID_METABOLISM_AND_TOXICITY-WP299
ELECTRON_TRANSPORT_CHAIN-WP111
IL12_AND_STAT4_DEPENDENT_SIGNALING_PATHWAY_IN_TH1_DEVELOPMENT
CYTOSKELETAL_REGULATION_BY_RHO_GTPASE
ANGIOGENESIS-WP1539
COMPLEMENT_ACTIVATION_CLASSICAL_PATHWAY-WP545
IL5_SIGNALING_PATHWAY
IL6_SIGNALING_PATHWAY
B_CELL_RECEPTOR_SIGNALING_PATHWAY-WP23
HISTAMINE_H1_RECEPTOR_MEDIATED_SIGNALING_PATHWAY
IL-1_P38
NES
logp

#### Supplementary Figure S3

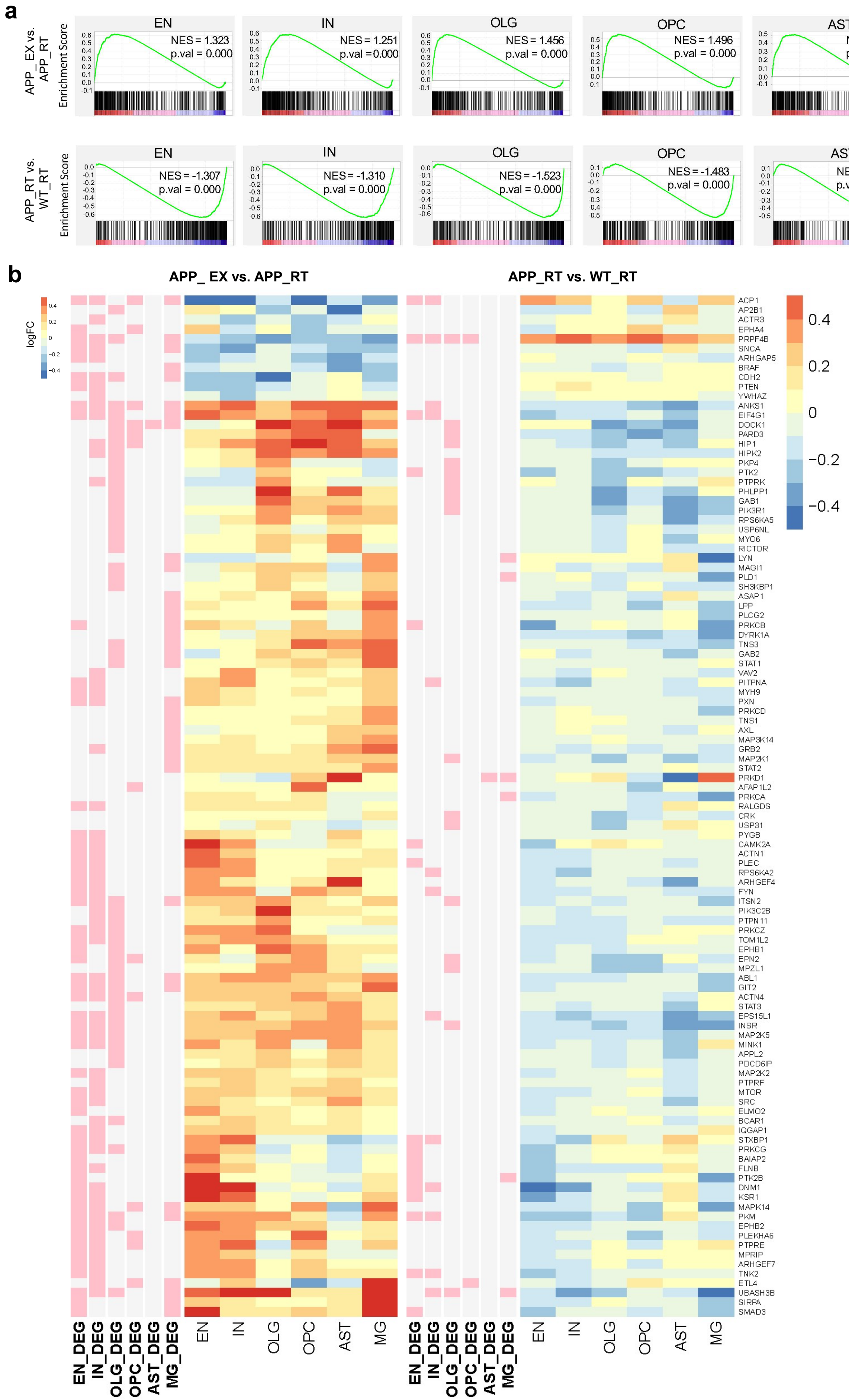

### Supplementary Figure S4

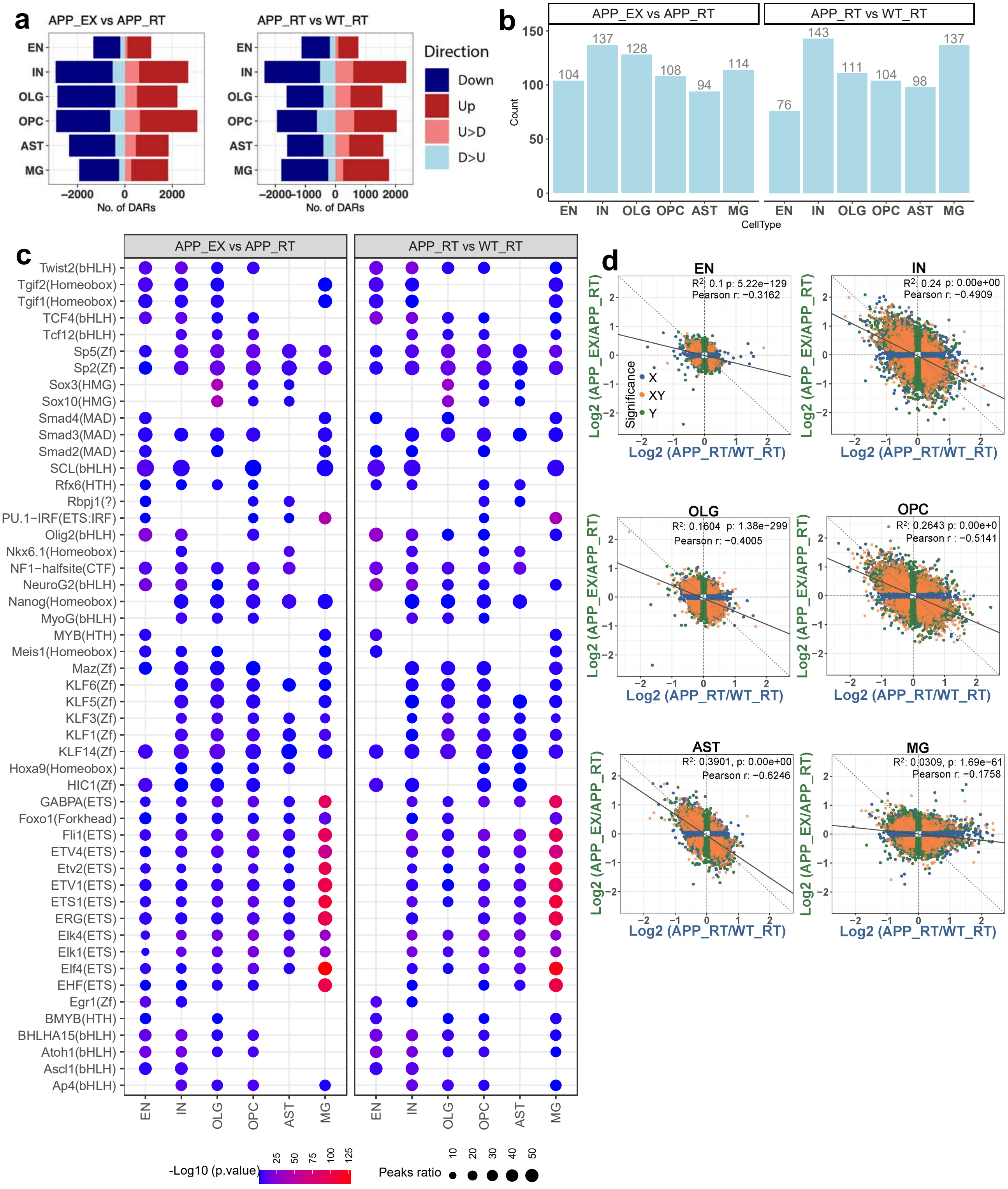

Supplmentary Figure S5

EN: APP_EX vs. APP_RT

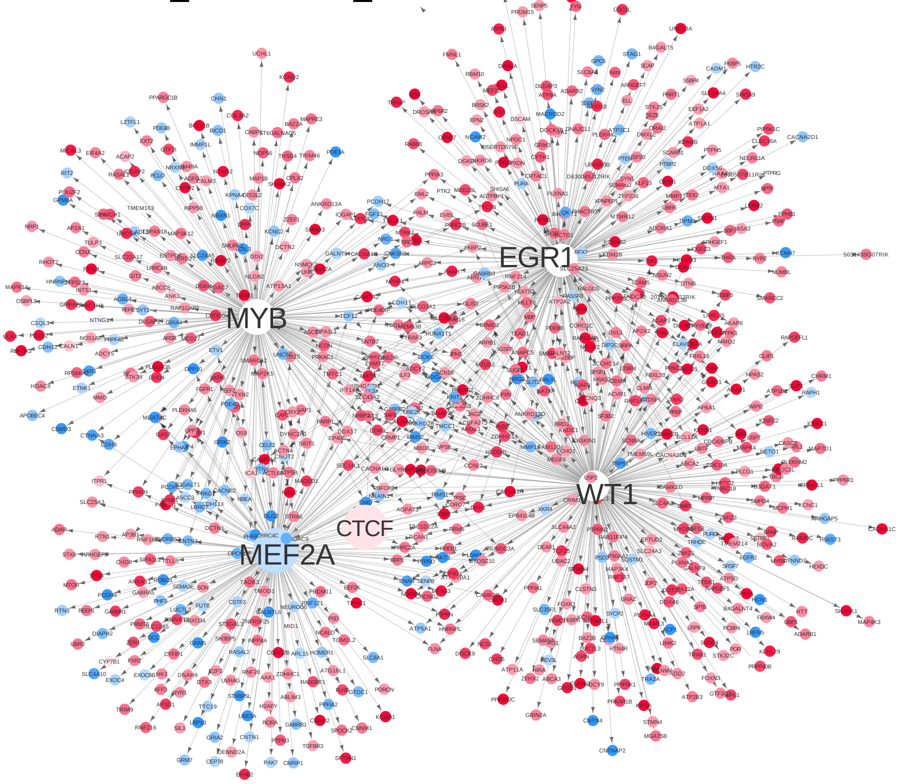


EN: APP_RT vs. WT_RT

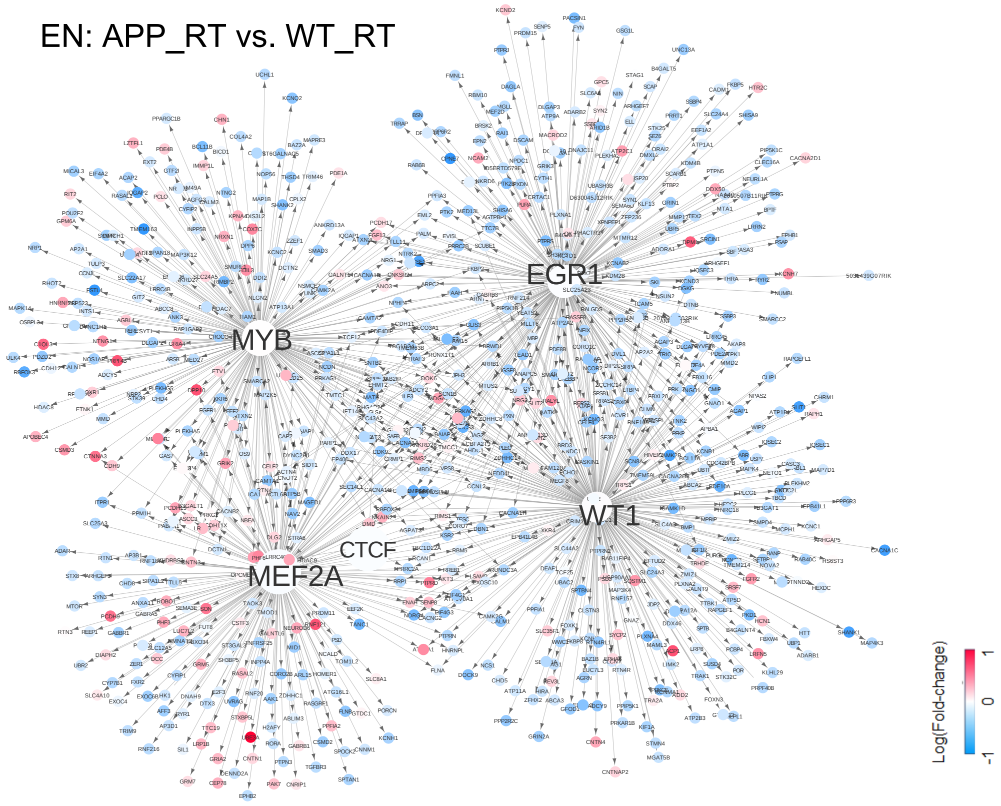

Supplementary Figure S6

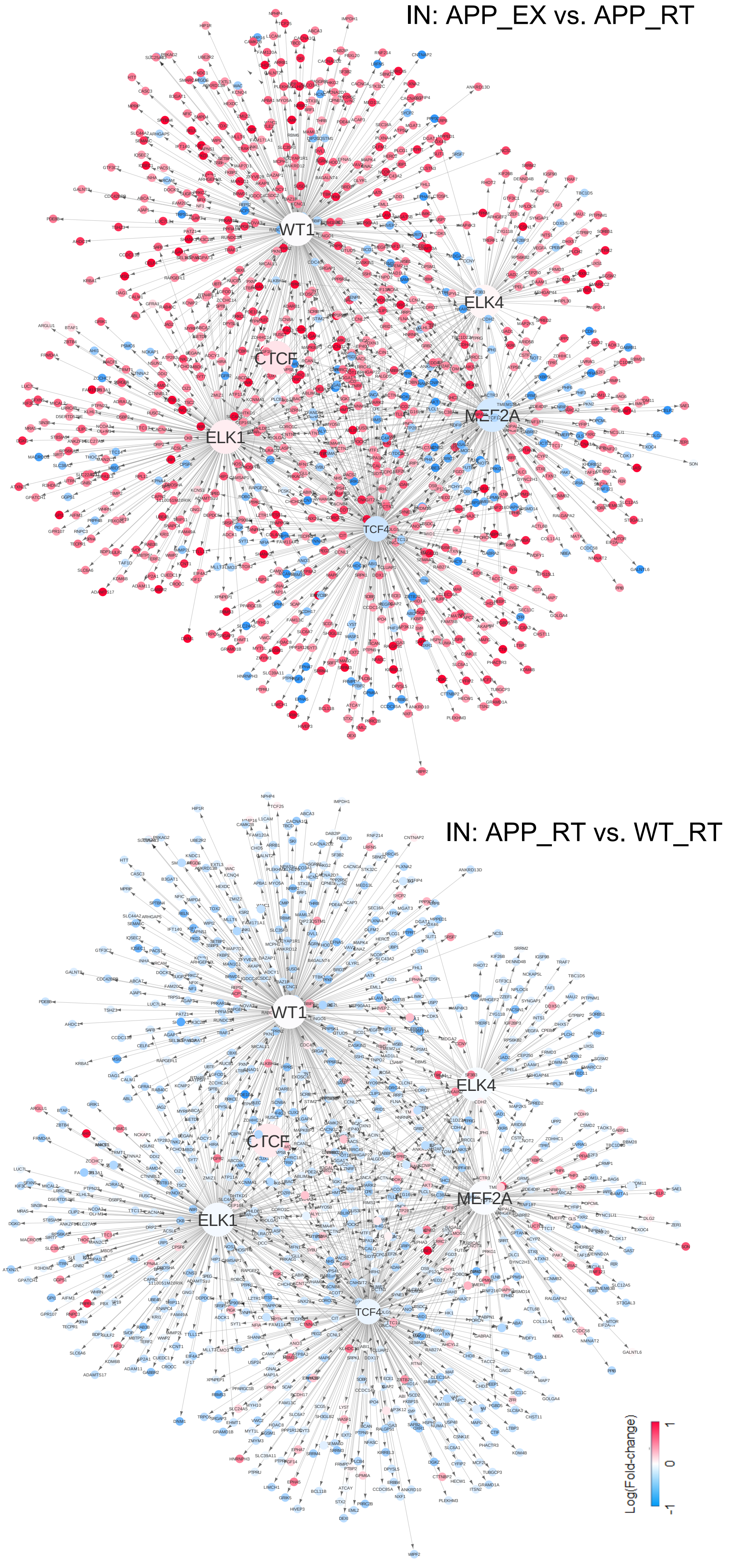

Supplementary Figure S7

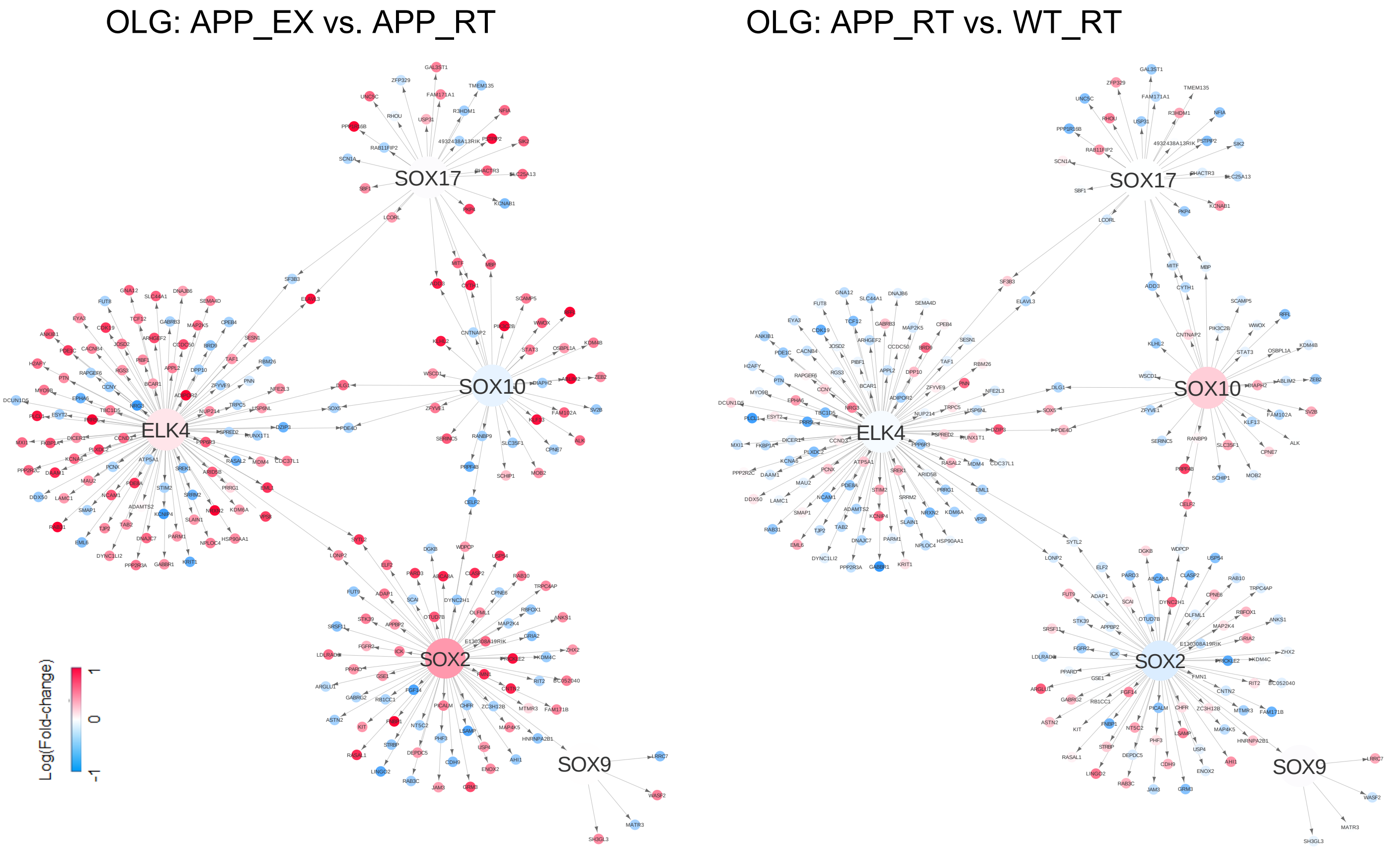

Supplementar Figure S8

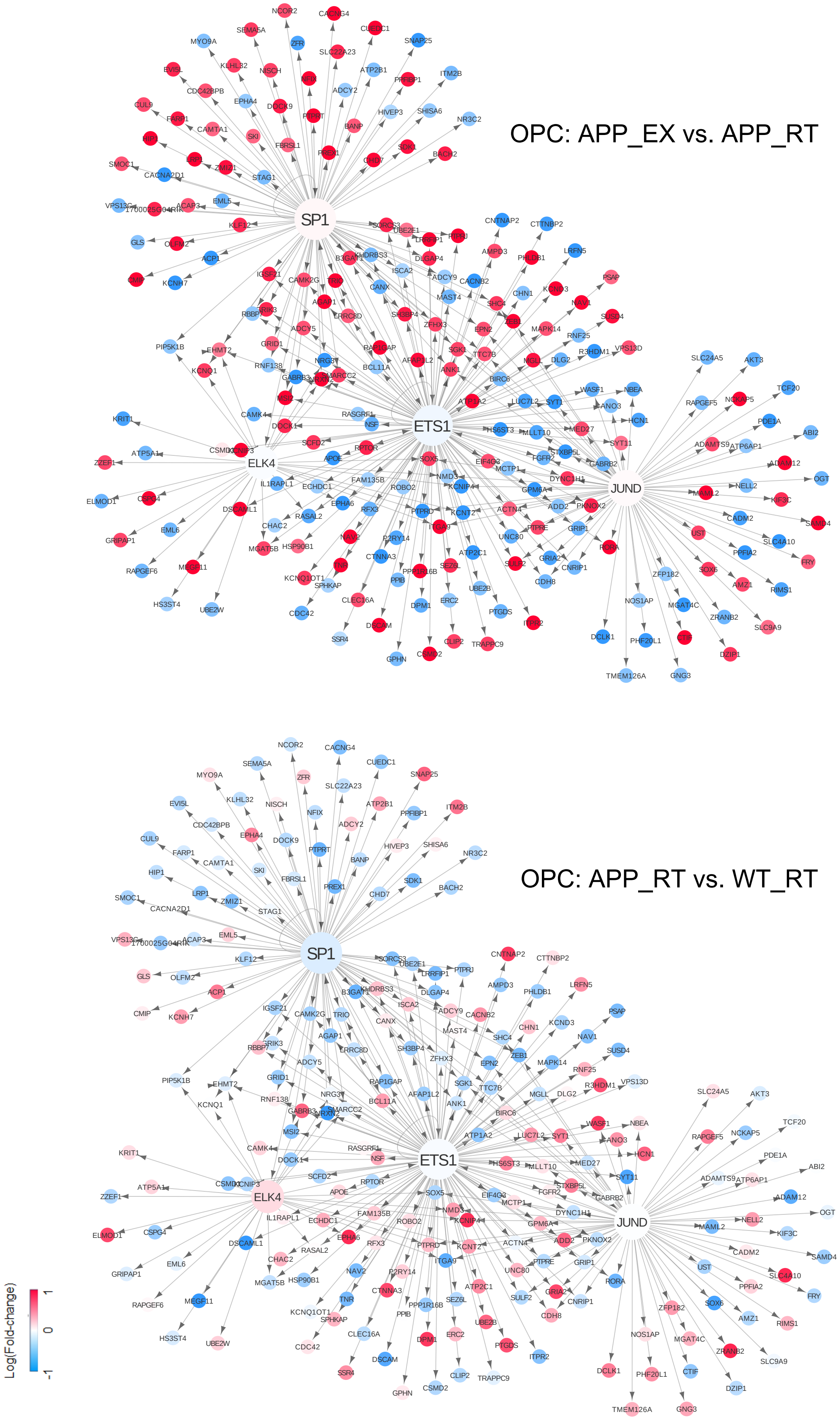

Supplementary Fig S9

#### AST: APP_EX vs. APP_RT

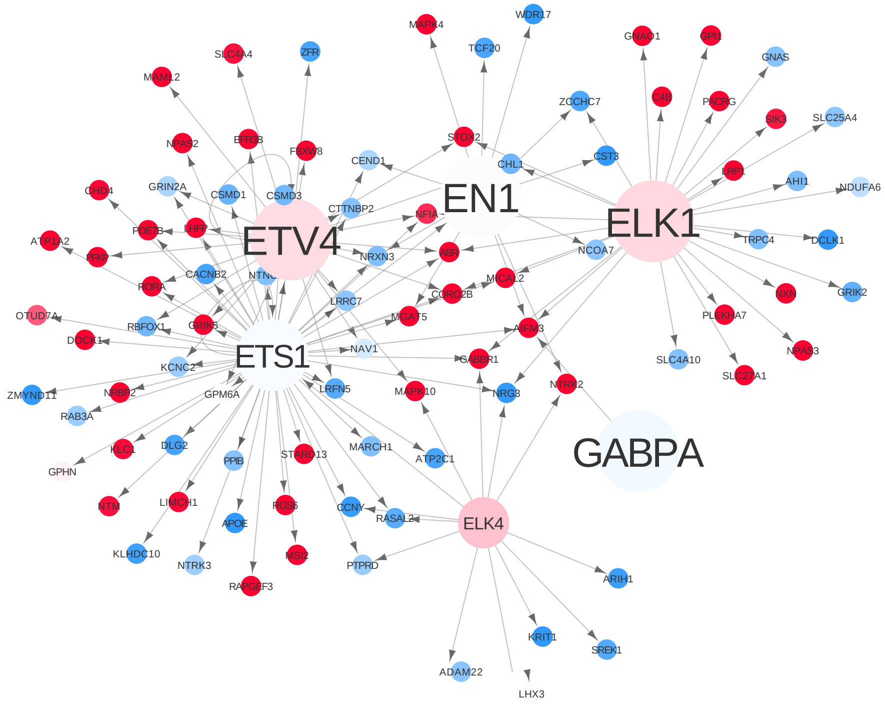


#### AST: APP_RT vs. WT_RT

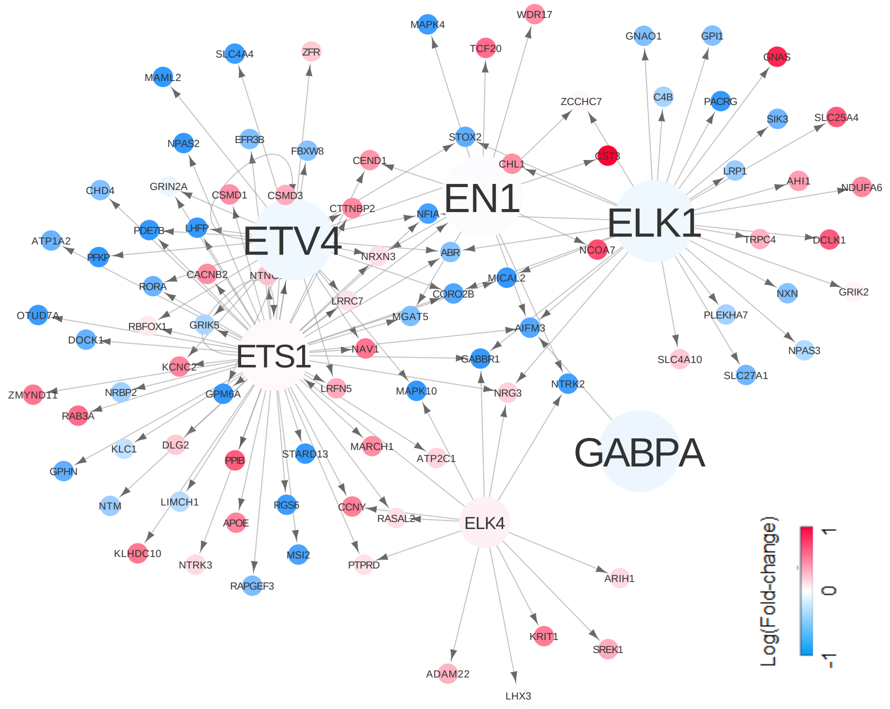

Supplementary Fig S10

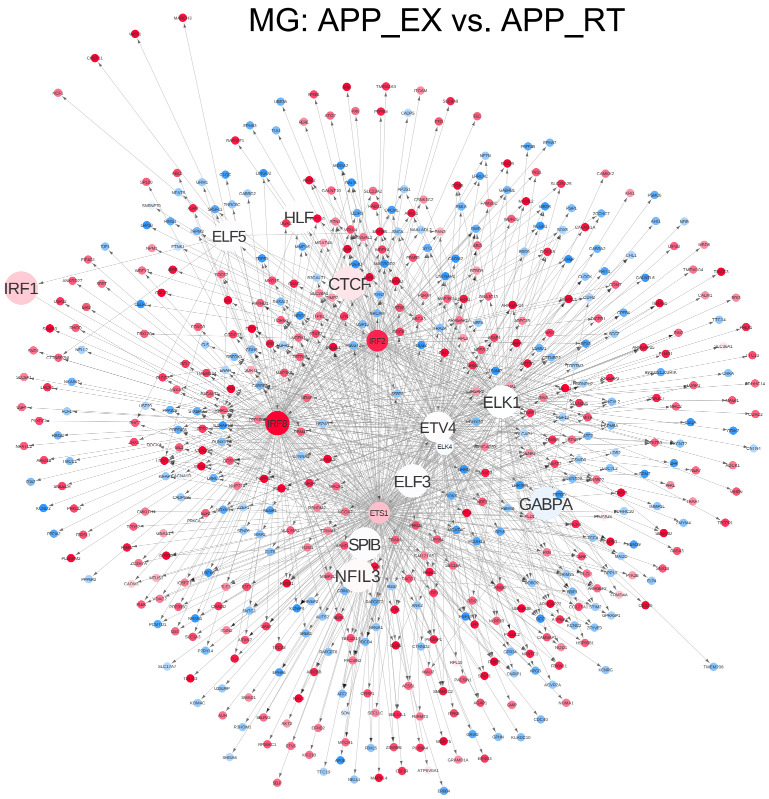


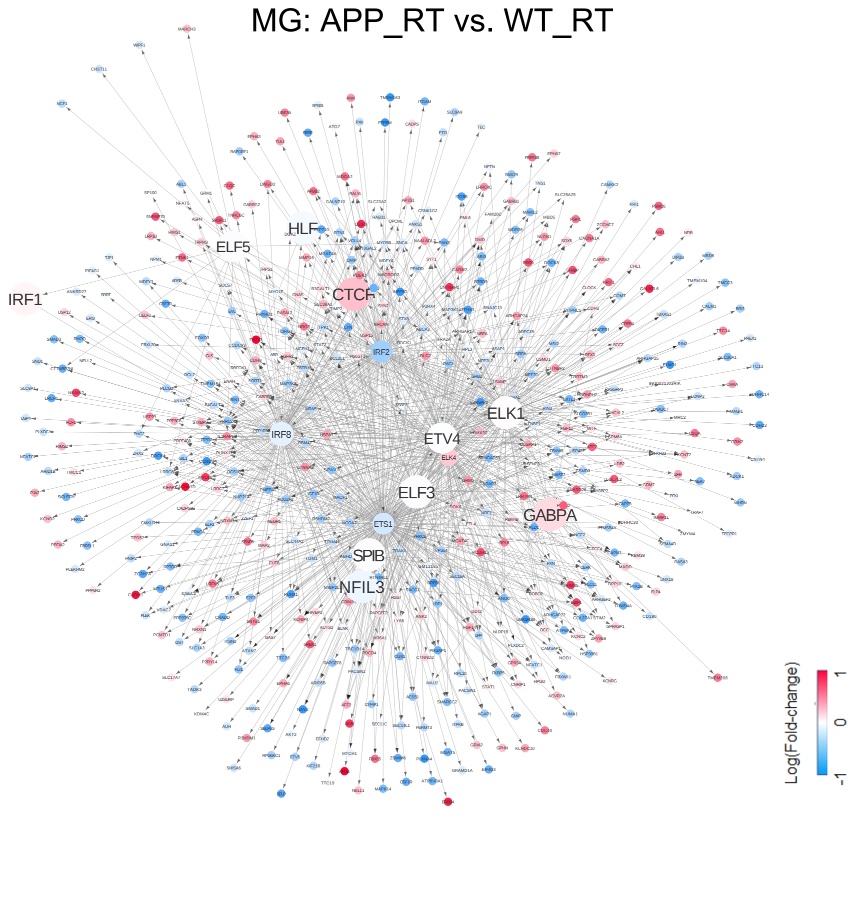

#### Supplementary Figure S11

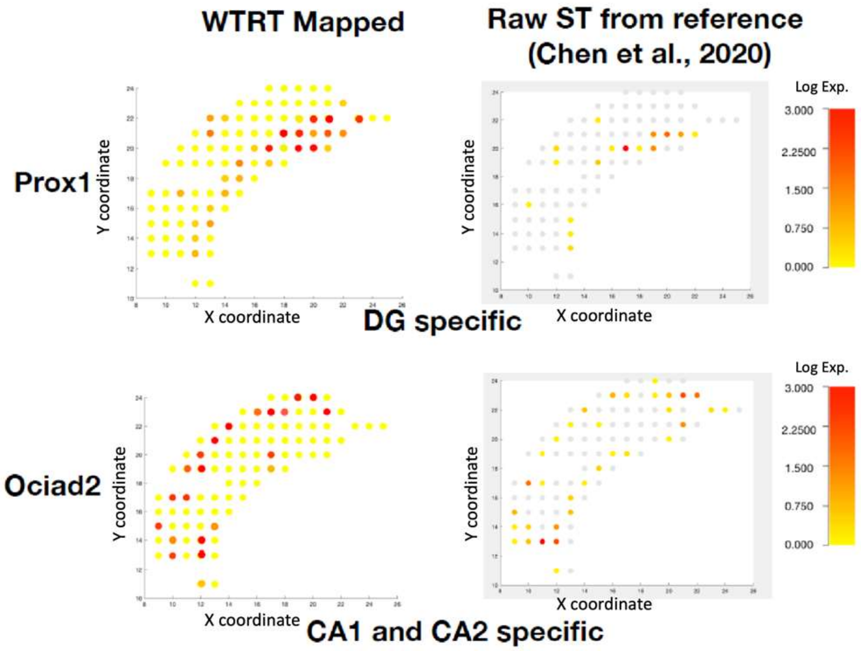
